# Circular Economy of Anaerobic Biofilm Microbiomes: A Meta-Analysis Framework for Re-exploration of Amplicon Sequencing Data

**DOI:** 10.1101/2020.12.23.424166

**Authors:** Ciara Keating, Anna Christine Trego, William Sloan, Vincent O’Flaherty, Umer Zeeshan Ijaz

**Author notes:** **Joint Corresponding Authors:** Dr Umer Z. Ijaz Prof. Vincent O’Flaherty, E E. Joint first authors, contributed equally to the study (Ciara Keating, Anna Christine Trego).

## Abstract

Use of high-throughput sequencing is widespread in efforts to understand the microbial communities in natural and engineered systems. Many built ecosystems, in particular those used for engineered wastewater treatment, have harnessed the metabolic capacity of complex microbial communities for the effective removal and recovery of organic pollutants. Recent efforts to better understand and precisely engineer such systems have increasingly used high-throughput sequencing to map the structure and function of wastewater treatment microbiomes. An enormous amount of data is readily available on online repositories such as the National Center for Biotechnology Information Short Read Archive (NCBI SRA). Here, we describe and provide an optimised meta-analysis workflow to utilise this resource to collate heterogenous studies together for anaerobic digestion research. We analysed 16S rRNA gene Illumina Miseq amplicon sequencing data from 31 anaerobic digestion studies (from high-rate digesters), including >1,300 samples. Additionally, we compare several methodological choices: extraction method, v-region, taxonomical database, and the classifier. We demonstrate that collation of data from multiple v-regions can be achieved by using only the taxa for which sequences are available in the reference databases, without losses in diversity trends. This is made possible by focusing on alternative strategies for taxonomic assignments, namely, bayesian lowest common ancestor (BLCA) algorithm which offers increased resolution to the traditional naïve bayesian classifier (NBC). While we demonstrate this using an anaerobic digestion wastewater treatment dataset, this methodology can be translated to perform meta-analysis on amplicon sequences in any field. These findings not only provide a roadmap for meta-analysis in any field, but additionally provide an opportunity to reuse extensive data resources to ultimately advance knowledge of wastewater treatment systems.

**Importance:** In this study, we have combined sequencing data from 31 individual studies with the purpose of identifying a meta-analysis workflow which can accurately collate data derived from sequencing different v-regions with minimal data loss and more accurate diversity patterns. While we have used Anaerobic Digestion (AD) communities for our proof-of-concept, our workflow (Fig 1) can be translated to any Illumina MiSeq meta-analysis study, in any field. Thereby, we provide the foundation for intensive data mining of existing amplicon sequencing resources. Such data-mining can provide a global perspective on complex microbial communities.

**Graphical Abstract:** 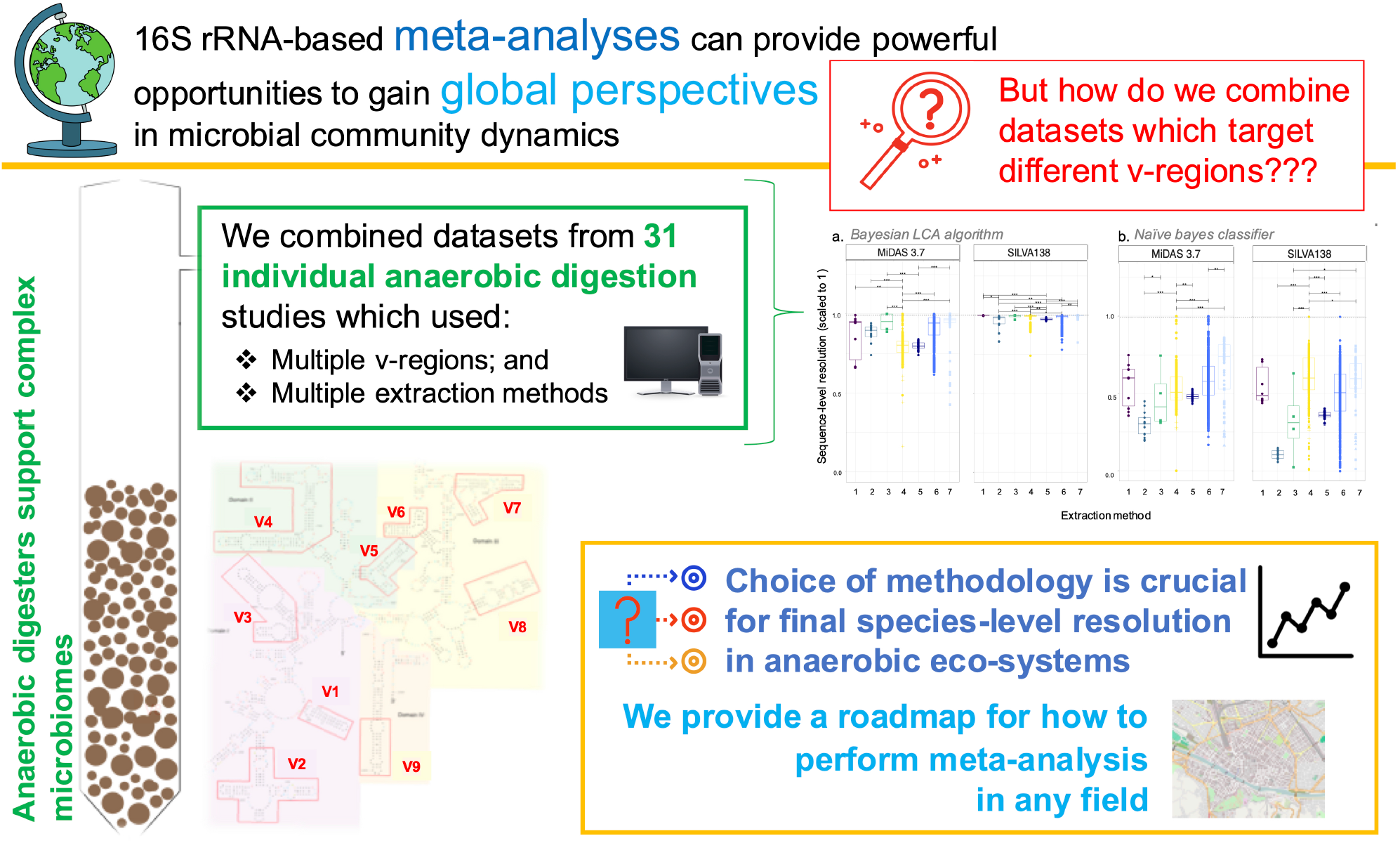

## Introduction

Complex microbial communities regulate global biogeochemical cycles and are exploited in engineered processes, such as biological wastewater treatment systems. Such systems, taking anaerobic digestion (AD) as an example are increasingly adopted for waste treatment and energy or intermediate platform chemical extraction [1]. For many years, molecular tools have been employed to gain a deeper understanding of the structure and function of these microbiomes. Methods include ‘old school’ molecular fingerprinting techniques [2] to more modern, short-read amplicon sequencing analyses or polymerase chain reaction (PCR)-free approaches such as metagenomics [3]. Amplicon sequencing has remained the popular and economic technique to understand the microbial community within environmental and engineered microbial systems. This has resulted in an unprecedented amount of data deposited in online repositories such as the National Center for Biotechnology Information (NCBI) short read archive (SRA) database.

**Figure 1.**
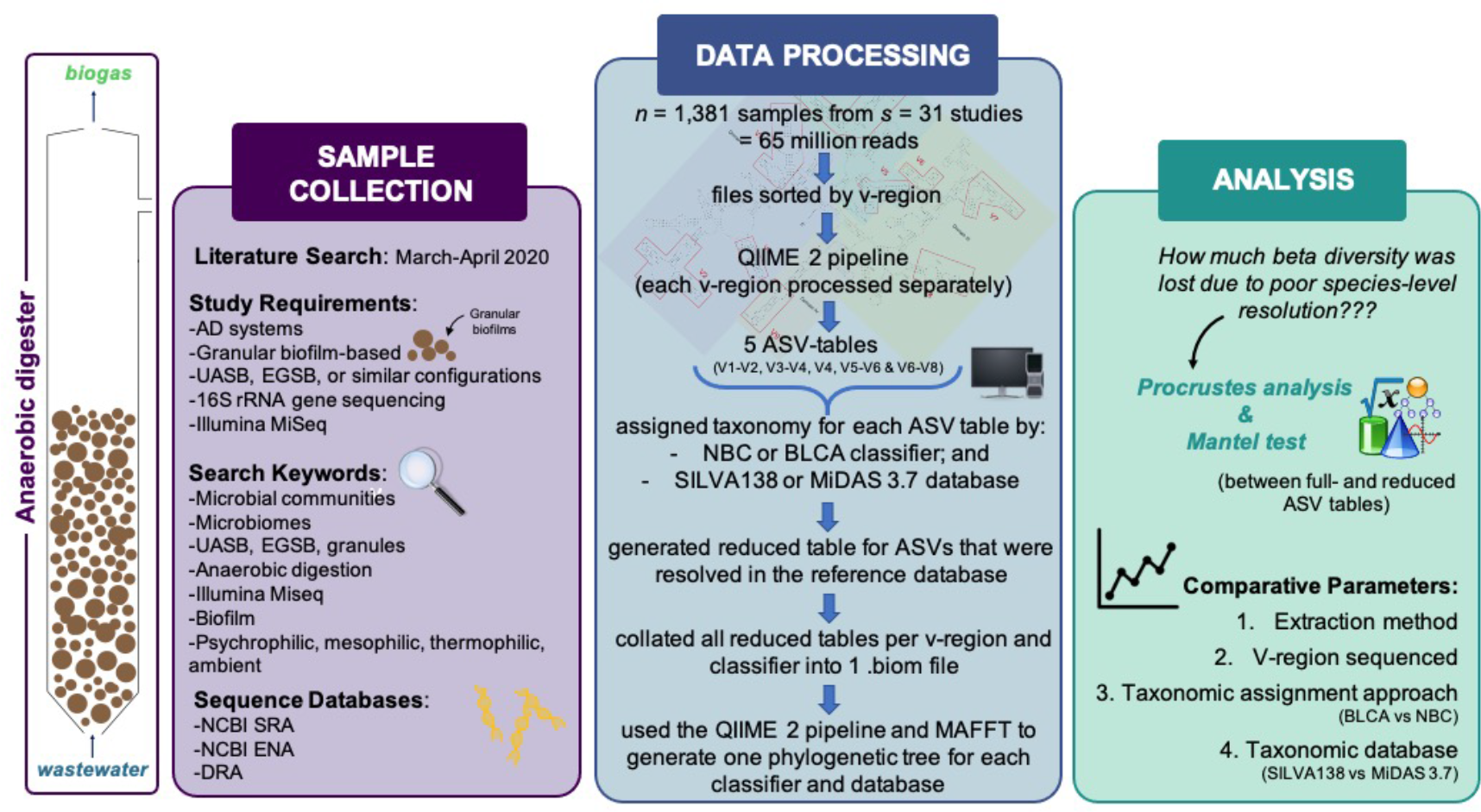
Overview of the meta-analysis workflow to assess the protocols used in the analysis of AD microbiomes.

There is a wealth of information held in this publicly available data that goes beyond the more specific questions addressed by any one study. By using and combining these previously published datasets (in a meta-analysis), we can not only discern large-scale patterns to provide a global perspective but can also interrogate our results in the context of other relevant works. Global analyses have been demonstrated as valuable in the understanding of oceanographic [4], activated sludge microbial ecosystems [5] and the ‘plastisphere’ [6] for example. In wastewater treatment systems we often describe ‘the circular economy’ and ‘resource recovery’ from waste. Thus, we propose to consider the data archived in online repositories not as a by-product of data accessibility or publishing but as a resource to be fully utilised later to further our understanding of complex microbial communities. However, there are hindrances. It is not possible to collectively analyse data arising from multiple variable regions (v-region). Indeed, many recent meta-analyses studies have incorporated only one v-region and therefore fail to provide a comprehensive perspective [7]. Alternatively, studies incorporating more than one v-region have been limited to taxonomic exploration as these studies have not developed a method for creating a collated phylogenetic tree [8]. To circumvent this problem, SATé-enabled phylogenetic placement (SEPP) technique has been proposed recently [9] which focuses on phylogenetic placement against a broad reference of phylogeny, and was then utilised to generate a collated phylogenetic tree in a meta-analysis study [6]. In our study, rather than considering optimal phylogenetic placement as above, we have focused on an alternative taxonomic approach to give reasonable resolution of species. A phylogenetic tree is necessary as it allows for a deeper understanding of the microbial community relationships and microbial community assembly processes [7], [10], [11]. In complex microbial systems such as wastewater treatment biofilms large-scale understanding of ecosystem function and taxa interaction through phylogenetic statistical analysis have the potential to unlock our ability to control or manipulate the community [12].

In terms of abundances, sophisticated approaches have appeared, such as k-d trees (k-dimensional tree) to combine multiple v-regions in terms of distances between the samples [13] on a landscape dictated by reference sequences. The limitations in all above approaches is the availability of fully-resolved sequences in the reference databases. Identification of key taxa, preferentially at the lowest taxonomic rank is required to obtain accurate information on important ecological traits [14] and species turnover [15]. Importantly, the archaea, which play a significant role in AD, are not well-represented in reference databases [16].

Another significant challenge is the large inconsistency of the methodologies applied in obtaining, preparing and analysing amplicon sequencing data. This is particularly true in the case of anaerobic digestion studies which lack a consensus set of protocols. DNA extraction method, library preparation (choice of v-region and primer sets), as well as post-hoc bioinformatic workflows crucially impinge on the final distribution of species taxonomy from complex environments [17]–[20]. Such distributions are important for cross-study comparisons and our overall understanding of AD process and function.

Cross-study comparisons can be limited due to the huge variation from extraction method right through to bioinformatic workflows [21]. Methodological biases are introduced at every step – from biomass sampling through to data processing [22]. In particular, the nucleic acid extraction method and choice of v-region (for 16S-based analysis) have been previously highlighted as critical choices [18], [19] in terms of DNA yields [23] and community diversity [24]. Popular extraction methods include variations of the phenol-chloroform co-extraction for DNA and RNA [25]. Or a plethora of kit-based DNA extraction options, among them, the FastDNA− Spin Kit for Soil (MP Biomedicals−, Burlingame, CA, USA) and the Powersoil® DNA Isolation Kit (MO BIO/QIAGEN, Carlsbad, CA, USA) are popular choices (Table 1). Generally, all methods will include a form of bead-beating for cell-lysis – the ferocity and duration of this particular step can vary across studies. The choice of v-region has a critical influence the final species distribution with primers varying in their coverage and showing bias towards specific phyla [26]. The V4 region is a popular choice, and universal primers, which simultaneously targeting bacteria and archaea (especially for wastewater-based studies). Though bacterial- and archaeal-specific primers are commonly used (Table 1).

**Table 1.**
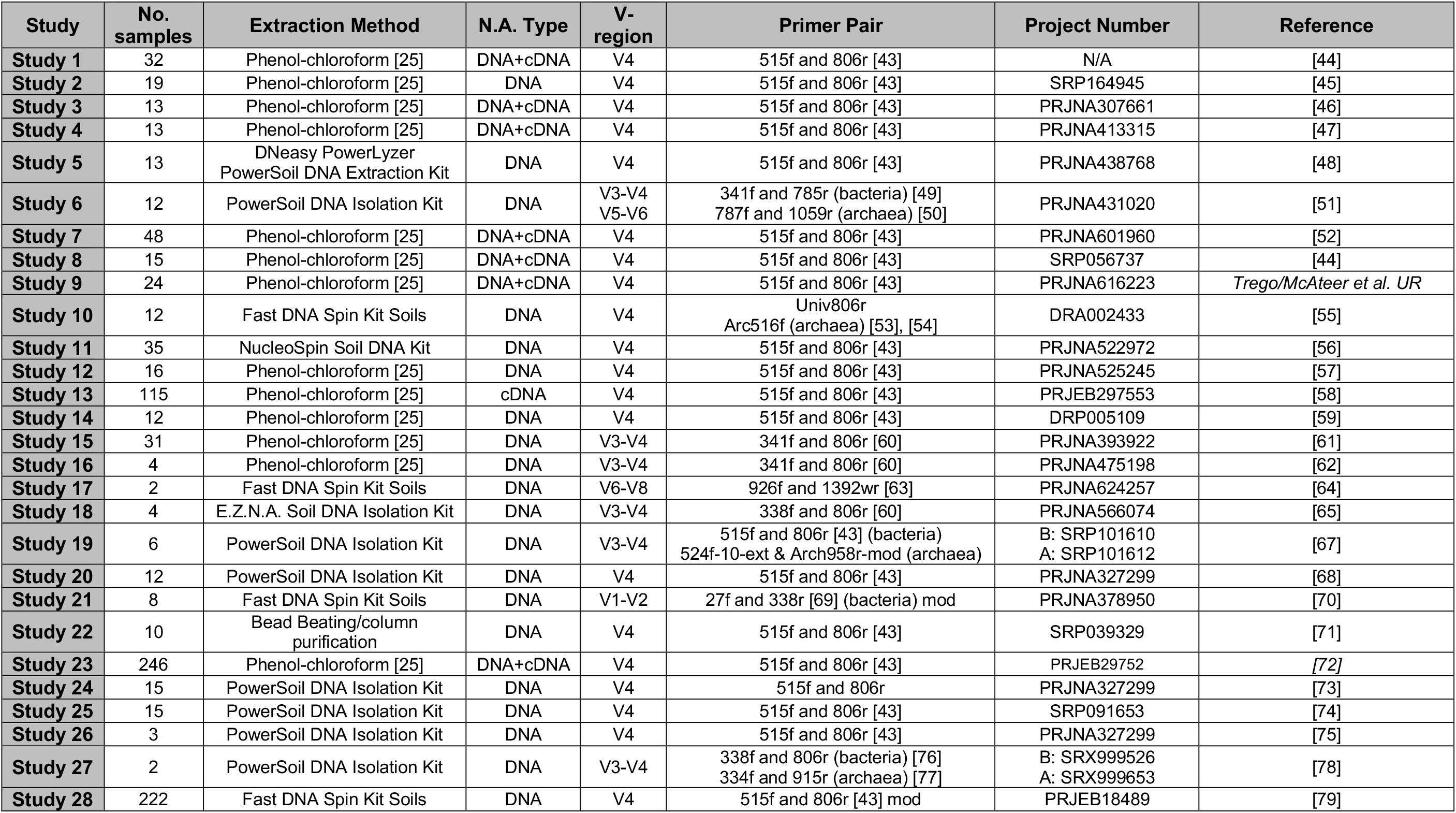
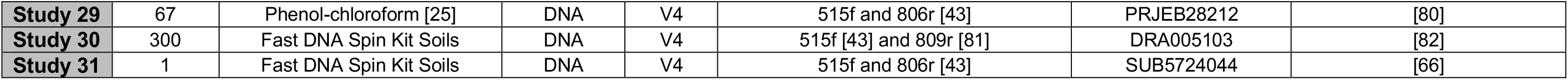
List of studies and relevant comparison parameters included in the meta-analysis.

Moreover, further variation may be introduced during post-hoc data processing [27]. Several processing pipelines have been developed with a series of steps and programs employed to align, denoise, and remove spurious sequences (i.e. MOTHUR [28], Qiime [29], KRAKEN [30]. While these pipelines are designed and updated for quality assurance, users typically follow a series of steps without deviation. Such pipelines typically resolve species through an operational taxonomic unit (OTU) approach [31], or an amplicon sequencing variant (ASV) approach [32] with the latter gaining popularity in recent years. In either an ASV or OTU-based approach, the sequences are aligned against a reference database to resolve their taxonomy up to species level, where possible. The alignment process is most often implemented as a classifier such as the traditional naïve bayesian classifier (NBC) [33]. Recently however, the bayesian lowest common ancestor (BLCA) algorithm [34] has been proposed to give shades-of-grey assignments (with confidences) by considering the phylogeny of the reference ‘hits’. This approach, as opposed to NBC, is able to provide greater species-level resolution [34]. In tandem, taxonomy is assigned using an online database. Currently, SILVA ribosomal RNA gene database (release 138) is the most comprehensive (containing 436,680 sequences) for 16S rRNA data [35]. However, species-level taxonomy is not curated within this database with species classifications often falling into the category of ‘uncultured’ or ‘metagenome’. In contrast, environment-specific and highly curated databases have also been developed, such as MiDAS [36] or AutoTax [14]. MiDAS focuses entirely on taxa from wastewater treatment systems with the MiDAS 3 database comprising 9,609 sequences derived from twenty activated sludge and AD systems in Denmark. It provides unique provisional names for unclassified microorganisms at species-level. Moreover, the MiDAS field guide provides functional and physiological data for unclassified organisms. In general, however, most AD studies continue to use broad databases such as SILVA 138 as standard within the chosen pipeline.

By implementing a meta-analysis approach, we can better understand how each methodological choice influences final species distributions and resolution. These types of analyses can therefore provide a roadmap for a set of design-rules for AD-microbiome research to yield greater species assignment and thus, great comparability across studies. Therefore, the aims of this study were two-fold: (i) to provide a roadmap for how to perform meta-analysis (i.e. the utility of combining data from various v-regions in view of taxonomy approaches); and (ii) to better understand how methodological choices impact final species-distributions in AD-microbiome analysis. To achieve this, we have comprehensively compared 31 previously published studies (>1,300 samples; Table 1) in terms of protocols applied (both wet-lab and bioinformatics workflows), to give an account of the differences resulting from the varied application of these approaches. We hypothesise that the choice of extraction protocol, amplicon v-region, taxonomy classification approach, and taxonomic database selection will have implications for the final taxa distribution and resolution. We then implemented a three step approach. First, sequences from each v-region were processed separately using either the SILVA138 or MiDAS 3 database and the NBC or BLCA methods of classification. Secondly, the resulting ASV-tables were collated only for those ASVs that were found in the reference database, and then finally, the full-length sequences of these ASVs were identified in the database to generate a consensus phylogenetic tree for further statistical analysis [37]–[40].

## Methods

### Study Selection

The studies selected as a part of this meta-analysis were chosen based on the following criteria: (i) AD communities sequenced from (ii) granular biofilms, or other biofilm samples, within (iii) high-rate upflow anaerobic bioreactors (UASB, EGSB or other similar configurations with retained biomass) sequenced using (iv) the Illumina MiSeq platform (Fig 1). Although there were several platforms from which to choose, here we have focused entirely on Illumina MiSeq datasets, which is the standard for most amplicon-based studies as the error rates are well-understood and are typically lower than other platforms [83]. Sequence files from the relevant samples (*n*=1,381 samples from *s*=31 previously published studies; Table 1) were downloaded using the SRA toolkit [84]. When studies did not provide a bioproject reference number, or neglected to upload the data, the corresponding authors were contacted to provide the raw sequence files.

Unfortunately, data accessibility was a significant challenge. 100+ studies with MiSeq datasets which could have been suitable for this meta-analysis study had yet to make their raw data publicly available (Supplementary File 1). Not only is this problematic in terms of reproducibility, but it hinders advancement of databases and puts limits on knowledge that can be gained from meta-analyses efforts.

### Meta-Analysis Workflow

The complete bioinformatics workflow, including linux codes and R scripts, is available (Supplementary File 2). The full workflow is highlighted in Figure 1. Briefly, datasets were categorised by v-region (V1-V2, V3-V4, V4, V5-V6 and V6-V8; Figure 1) and processed separately using the QIIME2 pipeline. Within QIIME2, the DADA2 denoising algorithm was used to obtain the ASVs (workflow provided at: https://github.com/umerijaz/tutorials/blob/master/qiime2_tutorial.md). This yielded 5 separate ASV tables – one for each v-region. Subsequently, taxonomy was assigned using one of two common approaches: the traditional naïve Bayesian classifier as part of the QIIME2 workflow [29], [33] and a Bayesian lowest common ancestor approach (BLCA; [34]. BLCA algorithm was ran according to the recommendations given at (https://github.com/qunfengdong/BLCA). Both taxonomic tools were used with either the wastewater treatment specific MiDAS 3.7 reference database (Nierychlo et al., 2020) or the broad SILVA 138 (Silva 138 SSURef Nr99) reference database. This resulted in 20 full ASV-tables (one per v-region; per classification method; per reference database). Reduced ASV-tables containing only the ASVs that were resolved through the reference database were retained. The ASV-tables were then collated to one biom file per classification method and reference database as described in Supplementary File 2 (four biom files in total). A consensus phylogenetic tree was then created using the reference sequences using the IDs from the reference database.

To determine whether collation at fully resolved ASVs (and therefore loss of ASVs that were not assigned) influenced our ability to corrupt diversity trends in the data, we performed Procrustes [85] and Mantel’s (1967) [86] statistical tests between Bray-Curtis dissimilarity measures from the original full ASV table and its reduced representation per v-region, classification method and reference database (Table 2). These tests measure the spatial distribution of a dataset [87]. Particularly, Procrustes rotates one dataset (full table) to maximum similarity with a target dataset (reduced table) by minimizing sum of square differences between them and then using correlation values as quality of fit. Mantel test, on the other hand is simply the correlation between entries of the two dissimilarity measures (full table and reduced table). In our results high correlation values (> 0.9) indicate minimal loss of beta diversity even with the removal of ASVs that are not resolved at sequence (species) level (Table 2).

**Table 2.**
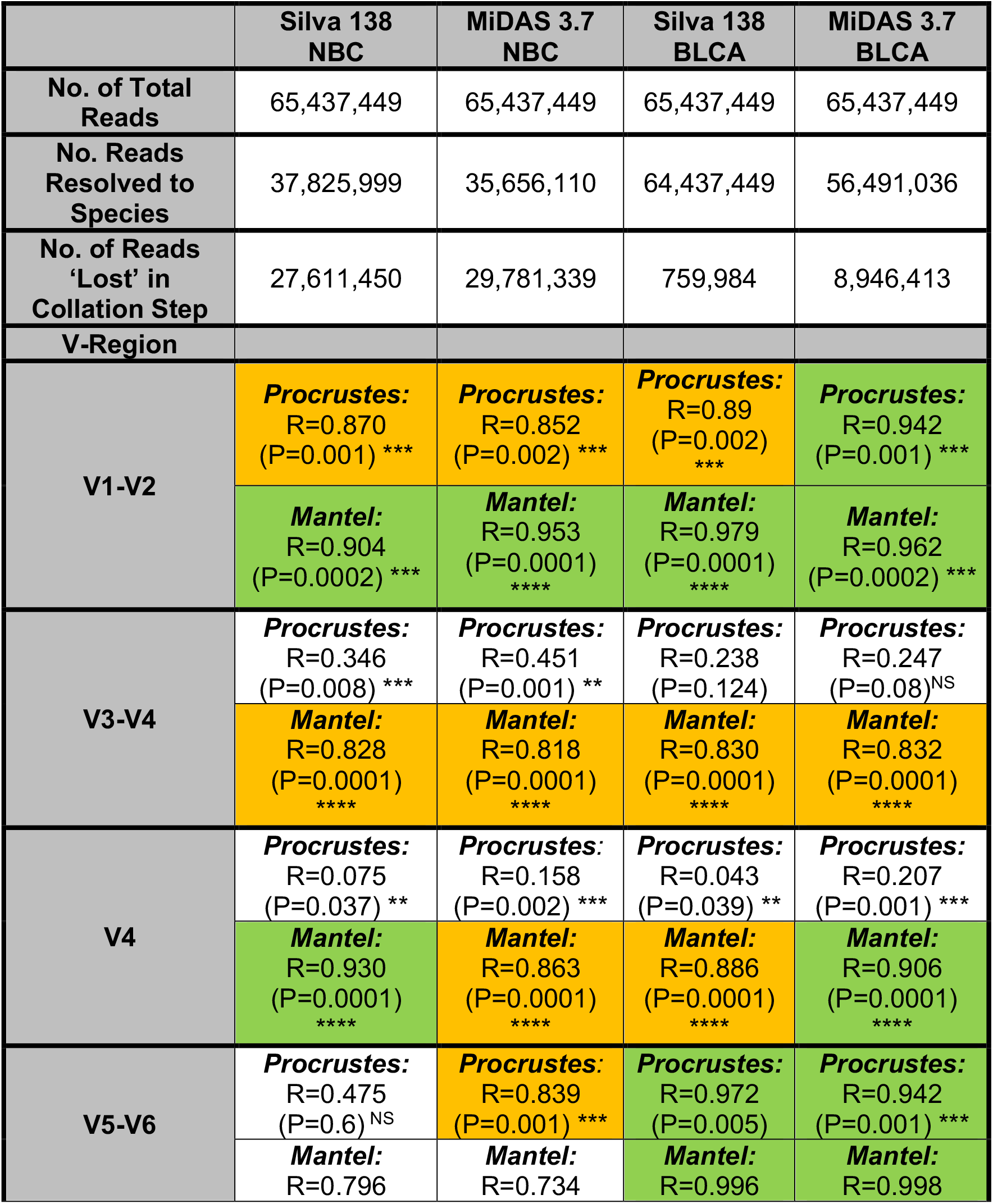
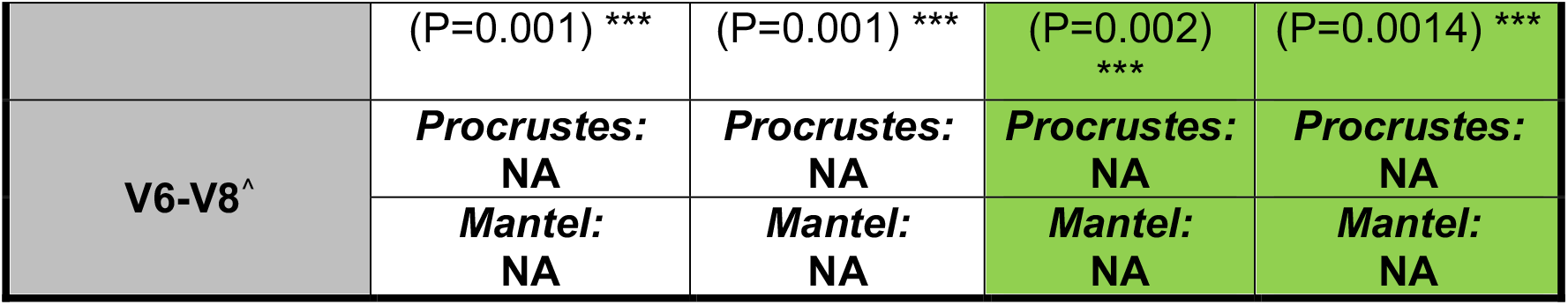
Summary of total reads and reads assigned to species-level using either the Silva 138 or MIDAS taxanomic database and the Bayesian Lowest Common Ancestor (BLCA) or Naïve Bayes Classifer (NBC) method’s of classification. Results of Procrustes and Mantel tests for each variable region (V1-V2, V3-V4, V4, V5-V6 and V6-V8^^^) using either the Silva 138 or MIDAS taxanomic database and the Bayesian Lowest Common Ancestor (BLCA) or Naïve Bayes Classifer (NBC) method’s of classification. Green colour indicates strong correlations R^2^ values > 0.9 and orange colour indicates reasonably strong correlations R^2^ values >0.8 which are in an acceptable range. P < 0.05*, P < 0.01**, P < 0.001***, P < 0.0001****. ^^^Only two samples so statistical analysis not available.

|  | Silva 138<br>NBC | MiDAS 3.7<br>NBC | Silva 138<br>BLCA | MiDAS 3.7<br>BLCA |
| --- | --- | --- | --- | --- |
| **No. of Total Reads** | 65,437,449 | 65,437,449 | 65,437,449 | 65,437,449 |
| **No. Reads Resolved to Species** | 37,825,999 | 35,656,110 | 64,437,449 | 56,491,036 |
| **No. of Reads 'Lost' in Collation Step** | 27,611,450 | 29,781,339 | 759,984 | 8,946,413 |
| **V-Region** |  |  |  |  |
| **V1-V2** | **Procrustes:**<br>R=0.870<br>(P=0.001) *** | **Procrustes:**<br>R=0.852<br>(P=0.002) *** | **Procrustes:**<br>R=0.89<br>(P=0.002) *** | **Procrustes:**<br>R=0.942<br>(P=0.001) *** |
|  | **Mantel:**<br>R=0.904<br>(P=0.0002) *** | **Mantel:**<br>R=0.953<br>(P=0.0001) **** | **Mantel:**<br>R=0.979<br>(P=0.0001) **** | **Mantel:**<br>R=0.962<br>(P=0.0002) *** |
| **V3-V4** | **Procrustes:**<br>R=0.346<br>(P=0.008) *** | **Procrustes:**<br>R=0.451<br>(P=0.001) ** | **Procrustes:**<br>R=0.238<br>(P=0.124) | **Procrustes:**<br>R=0.247<br>(P=0.08)<sup>NS</sup> |
|  | **Mantel:**<br>R=0.828<br>(P=0.0001) **** | **Mantel:**<br>R=0.818<br>(P=0.0001) **** | **Mantel:**<br>R=0.830<br>(P=0.0001) **** | **Mantel:**<br>R=0.832<br>(P=0.0001) **** |
| **V4** | **Procrustes:**<br>R=0.075<br>(P=0.037) ** | **Procrustes:**<br>R=0.158<br>(P=0.002) *** | **Procrustes:**<br>R=0.043<br>(P=0.039) ** | **Procrustes:**<br>R=0.207<br>(P=0.001) *** |
|  | **Mantel:**<br>R=0.930<br>(P=0.0001) **** | **Mantel:**<br>R=0.863<br>(P=0.0001) **** | **Mantel:**<br>R=0.886<br>(P=0.0001) **** | **Mantel:**<br>R=0.906<br>(P=0.0001) **** |
| **V5-V6** | **Procrustes:**<br>R=0.475<br>(P=0.6)<sup>NS</sup> | **Procrustes:**<br>R=0.839<br>(P=0.001) *** | **Procrustes:**<br>R=0.972<br>(P=0.005) | **Procrustes:**<br>R=0.942<br>(P=0.001) *** |
|  | **Mantel:**<br>R=0.796 | **Mantel:**<br>R=0.734 | **Mantel:**<br>R=0.996 | **Mantel:**<br>R=0.998 |
|  | (P=0.001) *** | (P=0.001) *** | (P=0.002) *** | (P=0.0014) *** |
| V6-V8^ | **Procrustes:**<br>NA | **Procrustes:**<br>NA | **Procrustes:**<br>NA | **Procrustes:**<br>NA |
|  | **Mantel:**<br>NA | **Mantel:**<br>NA | **Mantel:**<br>NA | **Mantel:**<br>NA |

## Results

### Collation at Database-resolved ASVs Conserves Trends in Beta-Diversity

In the collation step for the BLCA ASV-tables less than 14% of sequences were removed that did not identify to sequence (species) level. In contrast, with NBC as much as 45% of sequences were removed. To determine if this loss of sequences affects beta diversity of samples, we performed both Procrustes analyses and Mantel tests between the full ASV tables and the reduced ASV table (resolved to species-level). High correlations (high R^2^ values) indicated beta diversity dissimilarity was retained, irrespective of read loss. Correlation values were highly influenced by v-region, classification method and the reference database. The BLCA approach best-conserved beta diversity between full- and reduced-ASV tables (Table 2). This is evidenced by significantly higher correlations for most v-regions and both negligible read loss and diversity loss as compared to NBC (in both MiDAS 3.7 and SILVA138 reference databases). Notably, correlation values for V3-V4 were in the range of 0.8 (Mantel tests) indicating that the choice of v-region plays a vital role in terms of determining the final resolution.

### Extraction Method and V-region Impact Final Database-Resolution

The choice of both extraction method and v-region-amplified resulted in significant differences in sequence-level resolution (i.e. whether a database match was found; Fig 2). However, variation was observed in terms of classification method and database as well. The MiDAS3.7 + BLCA combination resulted in significant differences between extraction methods. In particular, the use of the (i) *DNeasy Powerlyzer Powersoil Extraction Kit*; (ii) *Fast DNA Spin Kit for Soil;* or the (iii) *NucleoSpin Soil DNA Kit* resulted in fewer matches to the database (Fig 2a). Conversely, the use of SILVA138 + BLCA resulted in very high sequence-level resolution (70-100% resolved) with less variability between methodological choices. Regardless of extraction method, the use of NBC resulted in lower sequence-level resolution with more variation within groups (Fig 2b). The NBC + SILVA138 combination, in particular, resulted in poorly-resolved sequences – especially when using the (i) *DNeasy Powerlyzer Powersoil Extraction Kit* (∼15% resolved); the (ii) *E*.*Z*.*N*.*A. Soil DNA Kit (∼40% resolved);* or the (iii) *NucleoSpin Soil DNA Kit (∼45% resolved)*. In terms of v-region, similar trends were observed. Notably, while the performance of most v-regions varied with respect to classification and database combination, V4 was consistently better. This indicates that when using regions other than V4, post-hoc data processing choices become critically important.

**Figure 2.**
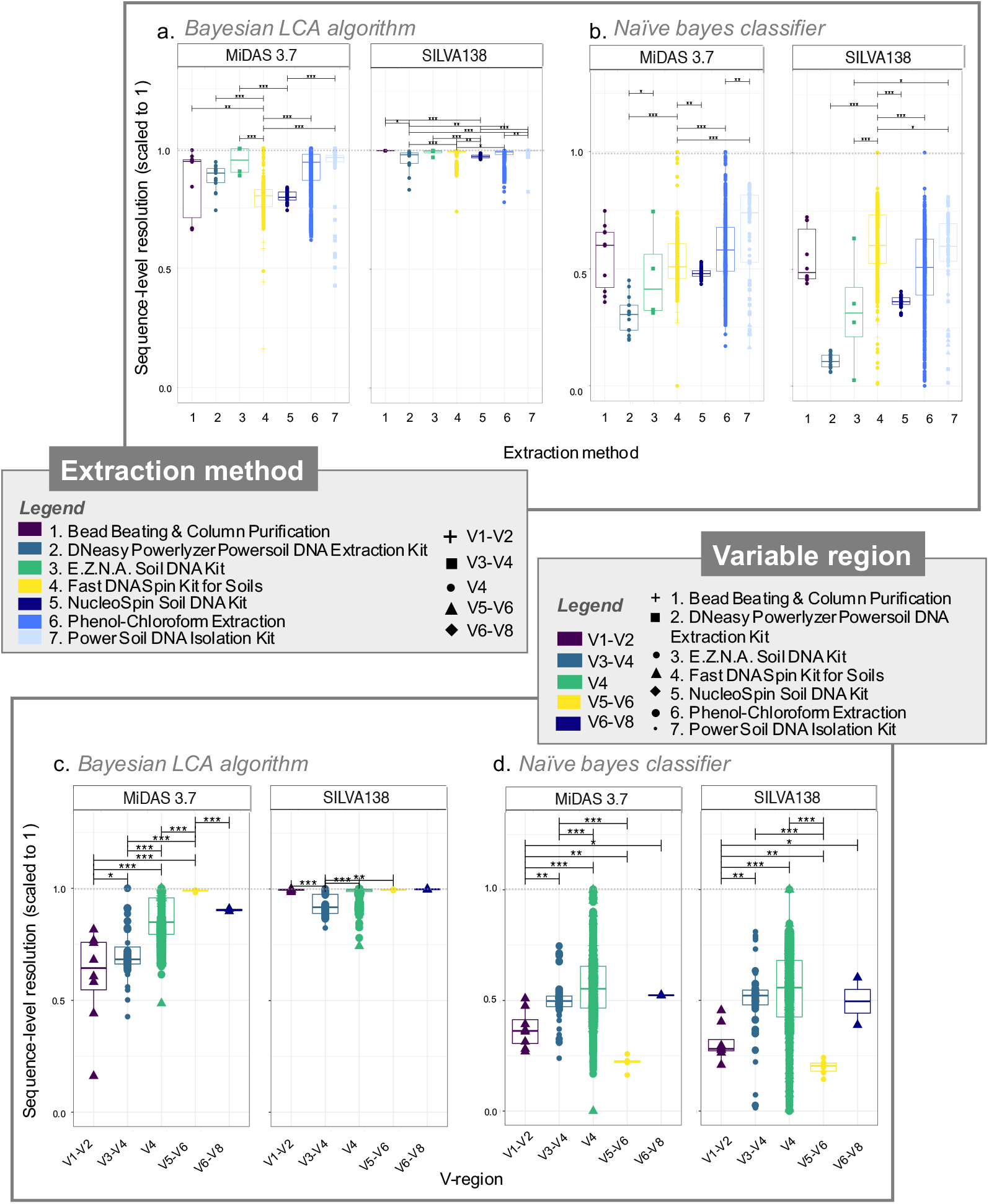
The number of ASVs resolved in the databases as a proportion of the total observed ASVs using both MiDAS 3.7 and SILVA138 databases for **(a)** BLCA and **(b)** NBC comparing different extraction methods; and **(c)** BLCA algorithm and **(d)** NBC comparing different hypervariable regions.

### BLCA yields greater sequence-level resolution than NBC

As previously noted, the BLCA approach not only resulted in the loss of fewer sequences (particularly in the case of SILVA 138), but better-conserved beta diversity than the traditional NBC approach (Table 2). Furthermore, the BLCA algorithm resulted in greater sequence-level resolution (Fig 2) and the use of BLCA, regardless of database selection or extraction method, resulted in fully resolved taxonomy (Fig 3) for both bacteria and archaea for the species-resolved reads. This means that from the proportion of sequences that were resolved, BLCA always provided information down to species-level (Rank 7).

**Figure 3.**
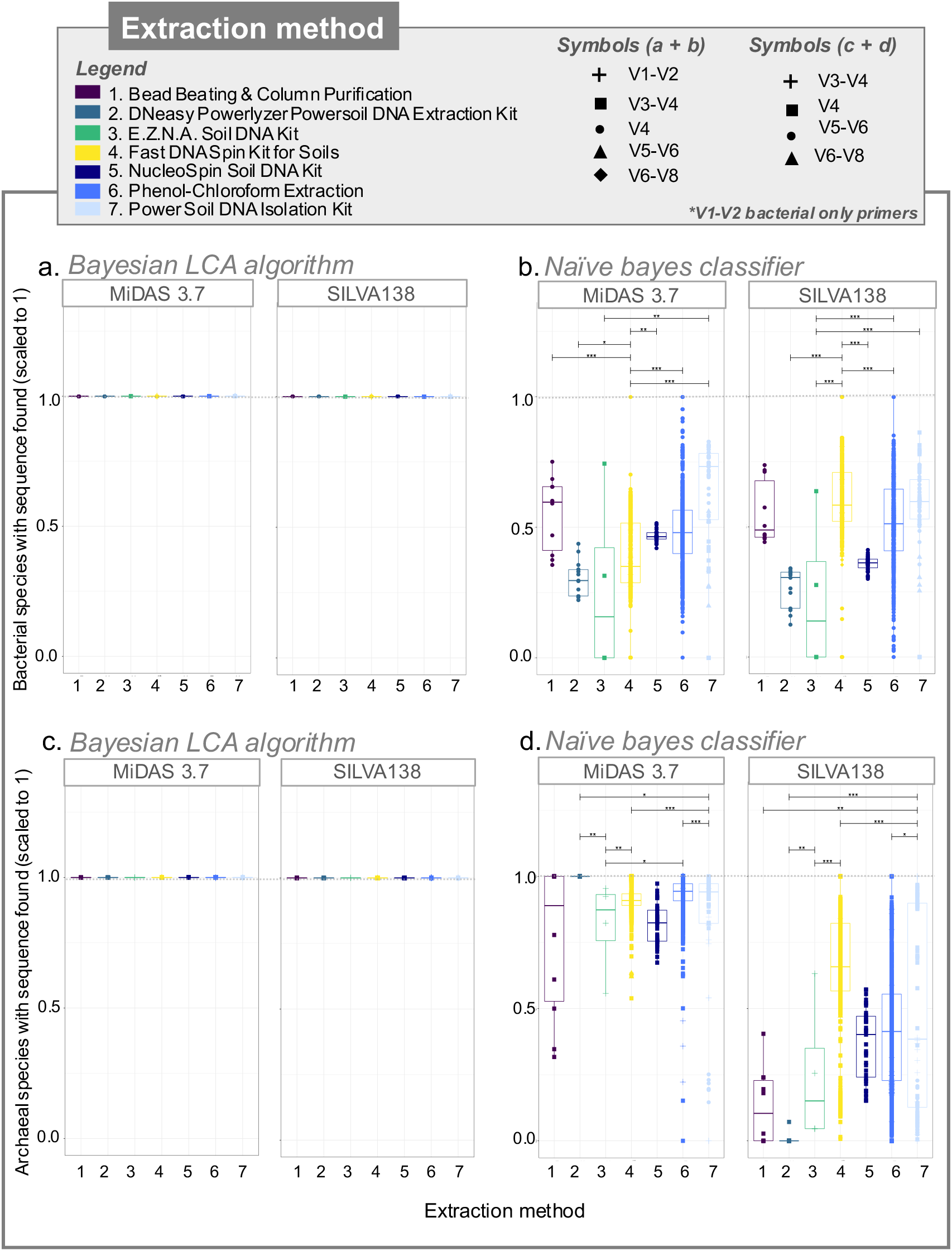
Bacterial and archaeal species-level resolution (i.e., all seven ranks were resolved) as a proportion of the total resolved ASVs (i.e., found in the reference database) with Superkingdom tag as “Bacteria/Archaea” using both MiDAS 3.7 and SILVA138 databases where **(a)** and **(c)** use BLCA for bacteria and archaeal groups, respectively; **(b)** and **(d)** use NBC for bacterial and archaeal groups, respectively all comparing different extraction methods.

### MiDAS database better identifies archaea – SILVA better identifies bacteria

Next, we interrogated how database and taxonomic algorithm choices separately affects bacterial and archaeal groups – both critical to the AD process. When using MiDAS + NBC we observed increased archaeal resolution (Fig 3d). With respect to bacteria, SILVA performed marginally better.

When we further interrogated the identity of the bacterial and archaeal species found in Figure 3, we observed that many returned incomplete nomenclature associated with uncultured species (Supplementary Figure S4). BLCA returned more ‘uncultured’ bacteria and archaea than NBC. Using the BLCA classifier approximately 70-90% of bacterial sequences found had incomplete taxonomic information. For the bacteria, the choice of reference database did not have an influence on the return of uncultured species. In contrast, in the archaeal assignments, Silva138 + BLCA provided poor taxonomic information with < 10% of sequences having complete taxonomy (90% ‘uncultured’). MiDAS, on the other hand, provided better archaeal taxonomic identification (10-60%). It must be noted that in the MiDAS database uncultured species are provided place-holder names in the form of midas_s_*x* which are highly curated.

## Discussion

### Meta-Analysis Workflow

One of the main aims of this study was to establish and examine a possible meta-analysis workflow to re-use amplicon sequencing data, in the context of wastewater treatment but broadly applicable to any other field. Since these studies targeted different variable regions within the 16S rRNA gene they are not comparable and therefore require separate processing through the bioinformatic workflow. Thus, an important aspect of meta-analysis is developing a robust approach to combine the output from a bioinformatic workflow. Collation is required for the separate ASV-tables which can be used to assess microbial taxa composition. To undertake more sophisticated statistical analysis a consensus phylogenetic tree is required (Lee et al., 2017; Stegen et al., 2012, 2013; Vass et al., 2020). Here we have demonstrated the utility of combining ASV-tables to only those ASVs that were fully resolved in the reference databases enabling us to generate a consensus phylogenetic tree using full-length sequences from the reference database. With phylogenetic tree available, recent and more powerful microbial community analyses are possible [38], [88]–[91]. For example, phylogenetic analysis can now provide information on community assembly mechanisms, diversity, and species recruitment, which are directly relevant to biofilm research. Our study clearly demonstrates that beta diversity can be conserved when combining data from multiple v-regions using appropriate taxonomy software. However, for the V3-V4 region, trends in beta-diversity were less well conserved. Thus, highlighting that data from the V3-V4 poses a challenge for meta-analyses work. It is anticipated that as databases continue to be curated, read-losses will decrease and this methodology of collation/meta-analysis will become even more robust. The valuable resource of gargantuan amplicon sequencing data in online repositories is a resource worth exploiting to understand complex microbiomes.

### Choice of methodology

Community analysis generally begins with the extraction of nucleic acids. The choice of methodology critically impacts final community distributions [19]. Here we show significant differences between 7 different extraction methods.

Differences were also dependent on downstream bioinformatics. For example, BLCA resulted in greater resolution for all extraction methods thereby overcoming pre-processing method variability. In particular, the combination of BLCA + SILVA138 provided high resolution and less variability between and within extraction methods. However, our study highlights that the phenol-chloroform co-extraction and Powersoil® DNA Isolation Kit were consistently among the best regardless of downstream processing. The phenol-chloroform method was the most popular from among the studies analysed here, likely due to accessibility and low-cost. Moreover, the method is designed to isolate both DNA and RNA simultaneously, giving access to active community profiles. High variability was observed using this method, likely due to user flexibility, particularly in terms of the addition of a bead-beating step for cell-lysis. In terms of extraction method choice, either of these methods are suitable.

The selection of primers and associated v-region is also an important choice. From among the 31 studies, the 515f and 806r universal primer pair amplifying the V4 region was the most popular choice (Table 1). While variability was observed in terms of downstream processing, overall, the V4 region seems to be a reasonable compromise, which was also previously noted [22]. AD studies could benefit from a standard primer set such as the standardisation efforts of the Earth Microbiome Project to allow for greater cross-study comparisons [92].

While extraction method and v-region did yield significant differences in terms of resolution of ASVs, taxonomic database and classification method had a much stronger impact. Taxonomic assignment tools have a clear effect on the underlying community resolution. BLCA had an outright advantage over the conventional method (NBC) by always maintaining a sequence match to the database in sequences that were resolved. This method significantly improves classification over existing methods as it calculates true sequence similarity between query sequences and database ‘hits’ using pairwise sequence alignment. Taxonomic assignments from species- to phylum-level are found according to a lowest common ancestor algorithm for each query sequence and are further assigned reliabilities through bootstrap confidence scores. This circumvents the limitations in the existing (NBC, etc.) assignment tools, which use probability-based criteria. NBC is however, widely adopted in traditional workflows (QIIME and MOTHUR).

However, we encountered a ‘bug’ whereby BLCA miss-identified genera (Rank 6), although the rest of the taxonomic path was correct, and this happened for only a handful of sequences. In light of this, we adopted a two-step approach (Supplementary File 2) where in the first step, we searched for the complete taxonomic path from phylum to species-level to identify which reference sequence the ASV is assigned to, and in the second pass, we searched for reference IDs based on last taxonomic level (Rank 7) to obtain the correct reference IDs. Regardless, BLCA seems to be the optimum choice for meta-analysis and for comprehensive taxonomic assignment.

Several general and environment-specific reference databases are available. For example, Genome Taxonomy Database Toolkit (GTDB-Tk - Chaumeil et al., 2019) and SILVA carry broad assignments for both bacteria and archaea, while databases such as MiDAS [36] and TaxAss (Rohwer et al., 2018) have been developed specifically for waste- and freshwater-related microbiomes respectively. MiDAS, in particular, is an ambitious effort to curate wastewater treatment microbiomes. We observed better archaeal species assignment with MiDAS. In contrast, SILVA138 provided in better bacterial assignment. As these databases continue to be updated with new sequences, MiDAS may gain an added advantage with respect to AD-microbiomes. The next anticipated release (MiDAS 4) will incorporate a global survey of anaerobic digesters and will likely increase in sequences relevant to biogas production. As such, one has to be judicious about finding a trade-off between bacterial and archaeal species-level resolution. This highlights the pressing need to have more archaeal sequences incorporated into broader databases given their importance in anaerobic systems or the supplementing of analysis with specific-databases. Not to mention the ongoing challenge with efforts to curate ‘uncultivated’ species [95]. The advantages of place-holder type strains in databases such as MiDAS [36] or AutoTAX [14] here would allow us to obtain greater insights into species turnover and function.

## Conclusion

In this study, we have combined sequencing data from 31 individual studies with the purpose of identifying a meta-analysis workflow which can accurately collate data derived from sequencing different v-regions with minimal data loss and more accurate diversity patterns. While we have used Anaerobic Digestion (AD) communities for our proof-of-concept, our workflow (Fig 1) can be translated to any Illumina MiSeq meta-analysis study, in any field. Thereby, we provide the foundation for intensive data mining of existing resources. In addition, due to the nature of our combined dataset, which incorporated >1,300 samples, we were able to provide AD-specific analysis on methodological choices from extraction method through to post-hoc bioinformatic workflows. Importantly, we highlight the influence of database and classification on archaeal assignments. Given the role of archaea in methane production in AD systems, a better resolution of key members of this kingdom is required.

## Author Contributions

UZI and VOF designed the study with input from CK and ACT. Data was collected by CK and ACT. Data was analysed by CK, ACT and UZI. ACT and CK prepared initial drafts of figures and drafted the paper and VOF, WS and UZI revised the document. All authors approve the paper and agree for accountability of the work therein.

## Supplementary Information

Supplementary information is available in a separate document for review.

## Acknowledgements

The authors thank all the authors of the included studies for providing their raw data and for responding to query requests. This work was financially supported by grants from the Higher Education Authority (HEA) of Ireland through: the Programme for Research at Third Level Institutions, Cycle 5 (PRTLI-5), co-funded by the European Regional Development Fund (ERDF); the Enterprise Ireland Technology Centres Programme (TC/2014/0016) and Science Foundation Ireland (14/IA/2371 and 16/RC/3889). This research was also funded by the Engineering and Physical Sciences Research Council (EPSRC) grants EP/K038885/1 and EP/P029329/1. UZI is funded by NERC Independent Research Fellowship NE/L011956/1.

