## Supplementary Information File 1 for "Circular Economy of Anaerobic Biofilm Microbiomes: A Meta-Analysis Framework for Re-exploration of Amplicon Sequencing Data"

| First Author | Year | Title | Journal | Platform |
|---|---|---|---|---|
| Studies W/O accession numbers from broad search of microbial communities in upflow anaerobic systems (March -April 2020) | | | | |
| Wang, W. | 2017 | Performance robustness of the UASB reactors treating saline phenolic wastewater and analysis of microbial community structure | Journal of Hazardous Materials | Illumina Miseq |
| Zhang, Z | 2018 | Enhancing anaerobic digestion and methane production of tetracycline wastewater in EGSB reactor with GAC/NZVI mediator | Water Reseach | Illumina Miseq |
| Li, Z. | 2019 | Performance and microbial community of an expanded granular sludge bed reactor in the treatment of cephalosporin wastewater | Bioresource Technology | Illumina Miseq |
| Zheng, Z | 2019 | Treatment of corn alcohol wastewater by anaerobic expanded granular sludge bed reactor and analysis of prokaryotic microbial community | Energy Sources, Part A: Recovery, Utiliz | Unspecified Ampli |
| Zhong, D | 2020 | Magnetite nanoparticles enhanced glucose anaerobic fermentation for bio-hydrogen production using an expanded granular sludge bed (EGSB) reactor | International Journal of Hydrogen Ener | Pyrosequencing |
| Lv, L | 2017 | Microbial community composition and function in a pilot-scale anaerobic-anoxic-aerobic combined process for the treatment of traditional Chinese medicine wastewater | Bioresource Technology | Illumina Miseq |
| Zhao, L | 2019 | Expanded granular sludge blanket reactor treatment of food waste at ambient temperature: Analysis of nitrogen compositions and microbial community structure | Bioresource Technology | Illumina Miseq |
| Xu, H. | 2018 | Granulation process in an expanded granular sludge blanket (EGSB) reactor for domestic sewage treatment: Impact of extracellular polymeric substances compositions and evolution of microbial populat | Bioresource Technology | Pyrosequencing |
| Yang, B | 2018 | Mechanism of high contaminant removal performance in the expanded granular sludge blanket (EGSB) reactor involved with granular activated carbon for low-strength wastewater treatment | Chemical Engineering Journal | Illumina Miseq |
| Miao, Y | 2017 | Assessment of phenol effect on microbial community structure and function in an anaerobic denitrifying process treating high concentration nitrate wastewater | Chemical Engineering Journal | Pyrosequencing |
| Fykse, E | 2016 | Microbial community structure in a full-scale anaerobic treatment plant during start-up and first year of operation revealed by high-throughput 16S rRNA gene amplicon sequencing | Bioresource Technology | Pyrosequencing |
| Ferrero, P | 2019 | Effect of substrate composition on the stability and microbial community of an anaerobic expanded granular sludge bed reactor treating printing solvent mixtures of ethanol and glycol ethers | International Biodeterioration & Biode | Illumina Miseq |
| Pan, X | 2019 | Impact of nano zero valent iron on tetracycline degradation and microbial community succession during anaerobic digestion | Chemical Engineering Journal | Illumina Miseq |
| Meng, L | 2017 | Amoxicillin effects on functional microbial community and spread of antibiotic resistance genes in amoxicillin manufacture wastewater treatment system | Journal of Environmental Sciences | Pyrosequencing |
| Liu, T | 2019 | Effect of influent pH on hydrolytic acidification performance and bacterial community structure in EGSB for pretreating crotonaldehyde manufacture wastewater after ozonation | Water Science & Technology | Illumina Miseq |
| Liao, R | 2018 | Temperature dependence of denitrification microbial communities and functional genes in an expanded granular sludge bed reactor treating nitrate-rich wastewater | RSC Advances | Pyrosequencing |
| Tian, T | 2018 | Bio-electrochemically assisting low-temperature anaerobic digestion of low-organic strength wastewater | Chemical Engineering Journal | Pyrosequencing |
| Tian, T | 2019 | Low-temperature anaerobic digestion enhanced by bioelectrochemical systems equipped with graphene/PPy- and MnO2 nanoparticles/PPy-modified electrodes | Chemosphere | Pyrosequencing |
| Cho, SK | 2018 | Effects of low-strength ultrasonication on dark fermentative hydrogen production: Start-up performance and microbial community analysis | Applied Energy | Ion PGM |
| Li, J | 2019 | Bacterial community structure and predicted function in an acidogenic sulfate-reducing reactor: Effect of organic carbon to sulfate ratios | Bioresource Technology | Illumina Miseq |
| Na, JG | 2016 | Microbial community analysis of anaerobic granules in phenol-degrading UASB by next generation sequencing | Biohemical Engineering Journal | Pyrosequencing |
| Li, H | 2018 | Performance, granule conductivity and microbial community analysis of upflow anaerobic sludge blanket (UASB) reactors from mesophilic to thermophilic operation | Biohemical Engineering Journal | Pyrosequencing |
| Zhuang, H | 2018 | Potential enhancement of direct interspecies electron transfer for anaerobic degradation of coal gasification wastewater using up-flow anaerobic sludge blanket (UASB) with nitrogen doped sewage slud | Bioresource Technology | Illumina Miseq |
| Wang, X | 2018 | Long-term effects of multi-walled carbon nanotubes on the performance and microbial community structures of an anaerobic granular sludge system | Applied Microbiology and Biotechnolog | Illumina Miseq |
| Lu, X | 2017 | Sulfidogenesis process to strengthen re-granulation for biodegradation of methanolic wastewater and microorganisms evolution in an UASB reactor | Water Reseach | Pyrosequencing |
| Xu, X | 2016 | Reductive Transformation of p-chloronitrobenzene in the upflow anaerobic sludge blanket reactor coupled with microbial electrolysis cell: performance and microbial community | Bioresource Technology | Pyrosequencing |
| Ou, C | 2016 | Coupling of iron shavings into the anaerobic system for enhanced 2,4-dinitroanisole reduction in wastewater | Water Reseach | Illumina Miseq |
| Li, Y | 2020 | Acidogenic and methanogenic properties of corn straw silage: Regulation and microbial analysis of two-phase anaerobic digestion | Bioresource Technology | Illumina Miseq |
| Niu, W | 2018 | Effect of fluctuating hydraulic retention time (HRT) on denitrification in the UASB reactors | Biochemical Engineering Journal | Illumina Miseq |
| Lee, J | 2017 | Bacteria and archaea communities in full-scale thermophilic and mesophilic anaerobic digesters treating food wastewater: Key process parameters and microbial indicators of process instability | Bioresource Technology | Pyrosequencing |
| Chen, C | 2017 | Evaluation of an up-flow anaerobic sludge bed (UASB) reactor containing diatomite and maifanite for the improved treatment of petroleum wastewater | Bioresource Technology | Illumina Miseq |
| Chen, C | 2019 | Characterization of aerobic granular sludge used for the treatment of petroleum wastewater | Bioresource Technology | Illumina Miseq |
| Lu, X | 2018 | Response of morphology and microbial community structure of granules to influent COD/SO42 – ratios in an upflow anaerobic sludge blanket (UASB) reactor treating starch wastewater | Bioresource Technology | Pyrosequencing |
| Torres, K | 2018 | Granulation and microbial community dynamics in the chitosan-supplemented anaerobic treatment of wastewater polluted with organic solvents | Water Reseach | Illumina Miseq |

| |
|---|
| Studies W/O accession numbers from keyword searching "Anaerobic Digestion" from Frontiers in Microbiology from the first 100 papers sorted by relevance - Conducted May 6, 2020 |
| none found - all had available accession numbers |

| | | | | |
|---|---|---|---|---|
| Studies W/O accession numbers from keyword searching "Anaerobic Digestion" from Water Research from the first 100 papers sorted by relevance - Conducted May 5, 2020 | | | | |
| Xiao, Y | 2019 | Autoinducer-2-mediated quorum sensing partially regulates the toxic shock response of anaerobic digestion | Water Research | Illumina Miseq |
| Wang, T | 2020 | Anaerobic digestion of sludge filtrate using anaerobic baffled reactor assisted by symbionts of short chain fatty acid-oxidation syntrophs and exoelectrogens: Pilot-scale verification | Water Research | Pyrosequencing |
| Li, Y | 2020 | High-efficiency methanogenesis via kitchen wastes served as ethanol source to establish direct interspecies electron transfer during anaerobic Co-digestion with waste activated sludge | Water Research | Illumina Hiseq |
| Zhao, Z | 2020 | Why do DIETers like drinking: Metagenomic analysis for methane and energy metabolism during anaerobic digestion with ethanol | Water Research | Ilumina Miseq |
| Zhang, W | 2020 | New insights into the effect of sludge proteins on the hydrophilic/hydrophobic properties that improve sludge dewaterability during anaerobic digestion | Water Research | Illumina Miseq |
| Wang, M | 2019 | Disposal of Fenton sludge with anaerobic digestion and the roles of humic acids involved in Fenton sludge | Water Research | Pyrosequencing |
| Yang, G | 2019 | Applying bio-electric field of microbial fuel cell-upflow anaerobic sludge blanket reactor catalyzed blast furnace dusting ash for promoting anaerobic digestion | Water Research | Undisclosed |
| Zhao, J | 2017 | Aged refuse enhances anaerobic digestion of waste activated sludge | Water Research | Illumina Miseq |
| Wu, L | 2016 | Long-term successional dynamics of microbial association networks in anaerobic digestion processes | Water Research | Illumina Miseq |
| Zhao, Z | 2017 | Towards engineering application: Potential mechanism for enhancing anaerobic digestion of complex organic waste with different types of conductive materials | Water Research | Pyrosequencing |
| Jang, HM | 2016 | Effect of increased load of high-strength food wastewater in thermophilic and mesophilic anaerobic co-digestion of waste activated sludge on bacterial community structure | Water Research | Pyrosequencing |
| Latif, M | 2017 | Influence of low pH on continuous anaerobic digestion of waste activated sludge | Water Research | pyrosequencing |
| Yang, Z | 2019 | Mitigation of ammonia inhibition through bioaugmentation with different microorganisms during anaerobic digestion: Selection of strains and reactor performance evaluation | Water Research | Illumina Miseq |

| Studies W/O accession numbers from keyword searching "Anaerobic Digestion" from Bioresource Technology from the first 100 papers sorted by relevance - Conducted May 6, 2020 | | | | |
|---|---|---|---|---|
| Navarro, R | 2020 | Combined simultaneous enzymatic saccharification and comminution (SESC) and anaerobic digestion for sustainable biomethane generation from wood lignocellulose and the biochemical characterizatic | Bioresource Technology | Illumina Miseq |
| Zhou, L | 2020 | Microbial community in in-situ waste sludge anaerobic digestion with alkalization for enhancement of nutrient recovery and energy generation | Bioresource Technology | Illumina Hiseq |
| Zhang, J | 2020 | Mixing strategies – Activated carbon nexus: Rapid start-up of thermophilic anaerobic digestion with the mesophilic anaerobic sludge as inoculum | Bioresource Technology | Illumina Hiseq |
| Wei, W | 2020 | Enhanced high-quality biomethane production from anaerobic digestion of primary sludge by corn stover biochar | BIoresource Technology | Illumina Miseq |
| Hülsen, T | 2020 | Anaerobic digestion of purple phototrophic bacteria – The release step of the partition-release-recover concept | Bioresource Technology | Illumina Miseq |
| Zhang, G | 2020 | Enhanced two-phase anaerobic digestion of waste-activated sludge by combining magnetite and zero-valent iron | Bioresource Technology | Pyrosequencing |
| Yin, F | 2020 | Additional function of pasteurisation pretreatment in combination with anaerobic digestion on antibiotic removal | Bioresource Technology | Undisclosed |
| Liu, Y | 2019 | Change to biogas production in solid-state anaerobic digestion using rice straw as substrates at different temperatures | Bioresource Technology | Pyrosequencing |
| Zhao, J | 2019 | Aged refuse enhances anaerobic fermentation of food waste to produce short-chain fatty acids | Bioresource Technology | Pyrosequencing |
| Xiao, Y | 2019 | Improved biogas production of dry anaerobic digestion of swine manure | Bioresource Technology | Pyrosequencing/Ill |
| Huang, H | 2020 | Sulfidogenic anaerobic digestion of sulfate-laden waste activated sludge: Evaluation on reactor performance and dynamics of microbial community | Bioresource Technology | Illumina Miseq |
| Xu, X | 2020 | Bioelectrochemical system for the enhancement of methane production by anaerobic digestion of alkaline pretreated sludge | Bioresource Technology | Pyrosequencing |
| Zhang, L | 2020 | Methane yield enhancement of mesophilic and thermophilic anaerobic co-digestion of algal biomass and food waste using algal biochar: Semi-continuous operation and microbial community analysis | Bioresource Technology | Pyrosequencing |
| Zhu, K | 2020 | Antagonistic effect of zinc oxide nanoparticle and surfactant on anaerobic digestion: Focusing on the microbial community changes and interactive mechanism | Bioresource Technology | Illumina Miseq |
| Dai, X | 2019 | Particle size reduction of rice straw enhances methane production under anaerobic digestion | Bioresource Technology | Illumina Miseq |
| Zhang, K | 2019 | Analysis for microbial denitrification and antibiotic resistance during anaerobic digestion of cattle manure containing antibiotic | Bioresource Technology | Illumina Hiseq |
| Wang, P | 2020 | Enhancement of biogas production from wastewater sludge via anaerobic digestion assisted with biochar amendment | Bioresource Technology | Undisclosed |
| Shi, X | 2020 | Two-stage anaerobic digestion of food waste coupled with in situ ammonia recovery using gas membrane absorption: Performance and microbial community | Bioresource Technology | Illumina Miseq |
| Wang, H | 2020 | Establishing practical strategies to run high loading corn stover anaerobic digestion: Methane production performance and microbial responses | Bioresource Technology | Undisclosed |
| Zhou, H | 2019 | Anaerobic digestion of aqueous phase from pyrolysis of biomass: Reducing toxicity and improving microbial tolerance | Bioresource Technology | Ion S5 |
| Sun, C | 2019 | Feasibility of dry anaerobic digestion of beer lees for methane production and biochar enhanced performance at mesophilic and thermophilic temperature | Bioresource Technology | Pyrosequencing |
| Zhang, J | 2019 | Anaerobic cultivation of waste activated sludge to inoculate solid state anaerobic co-digestion of agricultural wastes: Effects of different cultivated periods | Bioresource Technology | Illumina Miseq |
| Chen, H | 2020 | Dissecting methanogenesis for temperature-phased anaerobic digestion: Impact of temperature on community structure, correlation, and fate of methanogens | Bioresource Technology | Illumina Miseq |
| Suksong, W | 2020 | Biogas production from palm oil mill effluent and empty fruit bunches by coupled liquid and solid-state anaerobic digestion | Bioresource Technology | Illumina Miseq |
| Dalby, F | 2020 | Effect of tannic acid combined with fluoride and lignosulfonic acid on anaerobic digestion in the agricultural waste management chain | Bioresource Technology | Illumina Miseq |
| Wu, B | 2019 | Evaluating the effect of biochar on mesophilic anaerobic digestion of waste activated sludge and microbial diversity | Bioresource Technology | Illumina Miseq |
| Hu, Y | 2019 | Study of an enhanced dry anaerobic digestion of swine manure: Performance and microbial community property | Bioresource Technology | Illumina Hiseq |
| Xu, Y | 2020 | Enhancing methanogenesis from anaerobic digestion of propionate with addition of Fe oxides supported on conductive carbon cloth | Bioresource Technology | Undisclosed |
| Qin, Y | 2020 | Specific surface area and electron donating capacity determine biochar's role in methane production during anaerobic digestion | Bioresource Technology | Pyrosequencing/Ill |
| Zhang, Y | 2019 | Specific quorum sensing signal molecules inducing the social behaviors of microbial populations in anaerobic digestion | Bioresource Technology | Pyrosequencing/Ill |
| Wang, X | 2020 | Enhanced hydrolysis and acidification of cellulose at high loading for methane production via anaerobic digestion supplemented with high mobility nanobubble water | Bioresource Technology | Illumina Miseq |
| Ma, J | 2019 | Biochar triggering multipath methanogenesis and subdued propionic acid accumulation during semi-continuous anaerobic digestion | Bioresource Technology | Illumina Hiseq |
| Wang, G | 2020 | Redox-active biochar facilitates potential electron tranfer between syntrophic partners to enhance anaerobic digestion under high organic loading rate | Bioresource Technology | Illumina Miseq |
| Ma, J | 2019 | Effects of nano-zerovalent iron on antibiotic resistance genes during the anaerobic digestion of cattle manure | Bioresource Technology | Illumina Miseq |
| Salama, ES | 2020 | Enhanced anaerobic co-digestion of fat, oil, and grease by calcium addition: Boost of biomethane production and microbial community shift | Bioresource Technology | Illumina Miseq |
| Sun, W | 2019 | Solid-state anaerobic digestion facilitates the removal of antibiotic resistance genes and mobile genetic elements from cattle manure | Bioresource Technology | Illumina Hiseq |
| Bai, Y | 2019 | Sludge anaerobic digestion with high concentrations of tetracyclines and sulfonamides: Dynamics of microbial communities and change of antibiotic resistance genes | Bioresource Technology | Undisclosed |
| Feng, S | 2019 | Reutilization of high COD leachate via recirculation strategy for methane production in anaerobic digestion of municipal solid waste: Performance and dynamic of methanogen community | Bioresource Technology | Illumina Miseq |
| Zhi, S | 2019 | How methane yield, crucial parameters and microbial communities respond to the stimulating effect of antibiotics during high solid anaerobic digestion | Bioresource Technology | Illumina Miseq |

| | | | | |
|---|---|---|---|---|
| Studies W/O accession numbers from keyword searching "Anaerobic Digestion" from Water Science & Technology from the first 100 papers sorted by relevance - Conducted May 7, 2020 | | | | |
| Wang, J | 2019 | Enhancing anaerobic digestion of dairy and swine wastewater by adding trace elements: evaluation in batch and continuous experiments | Water Science & Technology | Illumina Hiseq |
| Hidaka, T | 2019 | Utilization of high solid waste activated sludge from small facilities by anaerobic digestion and application as fertilizer | Water Science & Technology | Illumina Miseq |
| Yanuka-Golu | 2019 | An electrode-assisted anaerobic digestion process for the production of high-quality biogas | Water Science & Technology | Illumina Miseq |
| Li, X | 2017 | An efficient method to improve the production of methane from anaerobic digestion of waste activated sludge | Water Science & Technology | Pyrosequencing |
| Park, S | 2016 | Mathematical models and bacterial communities for ammonia toxicity in mesophilic anaerobes not acclimated to high concentrations of ammonia | Water Science & Technology | Pyrosequencing |
| Studies W/O accession numbers from keyword searching "Anaerobic Digestion" from Environmental Science & Technology from the first 100 papers sorted by relevance - Conducted May 7, 2020 | | | | |
| Li, B | 2015 | Profile and Fate of Bacterial Pathogens in Sewage Treatment Plants Revealed by High-Throughput Metagenomic Approach | Environmental Science & Technology | Combination Hisec |
| Ma, L | 2017 | The Prevalence of Integrons as the Carrier of Antibiotic Resistance Genes in Natural and Man-Made Environments | Environmental Science & Technology | Illumina Hiseq |
| Xie, G | 2017 | Complete Nitrogen Removal from Synthetic Anaerobic Sludge Digestion Liquor through Integrating Anammox and Denitrifying Anaerobic Methane Oxidation in a Membrane Biofilm Reactor | Environmental Science & Technology | Pyrosequencing |
| Carey, D | 2016 | Triclocarban Influences Antibiotic Resistance and Alters Anaerobic Digester Microbial Community Structure | Environmental Science & Technology | Illumina Miseq |
| Duan, H | 2018 | Self-Sustained Nitrite Accumulation at Low pH Greatly Enhances Volatile Solids Destruction and Nitrogen Removal in Aerobic Sludge Digestion | Environmental Science & Technology | Illumina Miseq |
| Yang, Y | 2020 | Initial Deposition and Pioneering Colonization on Polymeric Membranes of Anaerobes Isolated from an Anaerobic Membrane Bioreactor (AnMBR) | Environmental Science & Technology | Illumina Miseq |
| Shi, X | 2015 | Investigation of Intertidal Wetland Sediment as a Novel Inoculation Source for Anaerobic Saline Wastewater Treatment | Environmental Science & Technology | Pyrosequencing |
| Studies W/O accession numbers from keyword searching "Anaerobic Digestion" from Applied and Environmental Science from the first 100 papers sorted by relevance - Conducted May 8, 2020 | | | | |
| Dang, P | 2013 | Methanosarcinaceae and Acetate-Oxidizing Pathways Dominate in High-Rate Thermophilic Anaerobic Digestion of Waste-Activated Sludge | Applied and Environmental Microbiolo | Pyrosequencing |
| Zhou, L | 2015 | Transcriptomic and Physiological Insights into the Robustness of Long Filamentous Cells of Methanosaeta harundinacea, Prevalent in Upflow Anaerobic Sludge Blanket Granules | Applied and Environmental Microbiolo | Ion torrent |
| Studies W/O accession numbers from keyword searching "Anaerobic Digestion" from FEMS Ecology from the first 100 papers sorted by relevance - Conducted May 8, 2020 | | | | |
| Kümmel, S | 2015 | Anaerobic naphthalene degradation by sulfate-reducing Desulfobacteraceae from various anoxic aquifers | FEMS Ecology | Illumina Miseq |
