## Supplementary Information File 2 for "Circular Economy of Anaerobic Biofilm Microbiomes: A Meta-Analysis Framework for Re-exploration of Amplicon Sequencing Data"

**Supplemental File 2 (Meta-analysis workflow codes)**

### Downloading data using SRI Toolkit

Go to BioProject website and search for a particular keyword (for example, “anaerobic digestion”). Download SRA toolkit from: https://www.ncbi.nlm.nih.gov/sra and use the following one-liners on linux terminal to get the paired-end files for a relevant bioproject.

Step 1: Get all the SRR numbers associated with a bioproject PRJNA*

```
esearch -db sra -query PRJNA302804 | efetch --format runinfo |cut -d
"," -f 1 > SRR.numbers
```

Step 2: Retain only the SRR numbers in the files

```
awk '/SRR/' SRR.numbers > SRR.numbers.filtered
```

You can either use a single SRR number (SRR3233069) with fastq-dump to download the corresponding files

```
fastq-dump --split-files --origfmt --gzip SRR3233069
```

of you can use the command in a for loop

```
for i in $(cat SRR.numbers.filtered); do echo Processing $i; fastq-
dump --split-files --origfmt --gzip $i ; done
```

<u>Sequences were sorted into folders per variable region. The below pipeline was then carried out per v-region.</u>

### QIIME2 workflow

**Step 1**: We are going to organize our data in such a manner that for every sample we have the folder name extracted from the paired-end files, and we are going to dump the raw sequences in a “Raw” folder:

```
[User@HPC ~/user/sequences]$ for i in $(awk -F"_" '{print $1}' <(ls
*.fastq) | sort | uniq); do mkdir $i; mkdir $i/Raw; mv $i*.fastq
$i/Raw/.; done

[User@HPC ~/user/sequences]$ cd ..
```

```
[User@HPC ~/user]$ mkdir qiime2_tutorial
[User@HPC ~/user]$ cd qiime2_tutorial
[User@HPC ~/user/qiime2_tutorial]$
```

There are different ways in which we can import data to qiime2:
https://docs.qiime2.org/2020.2/tutorials/importing/

```
[User@HPC ~/user/sequences]$ d="/home/eng/User/user/sequences/";
[User@HPC ~/user/sequences]$ cd ../qiime2_tutorial
```

Next step is to generate fictitious barcodes required to import data in Earth Microbiome Project (EMP) format (consult http://userweb.eng.gla.ac.uk/user.ijaz/bioinformatics/oneliners.html#PERLOL on how I have written one-liner in perl to generate fictitious barcodes):

```
[User@HPC ~/user/qiime2_tutorial]$ t=$(ls $d | wc -l);
[User@HPC ~/user/qiime2_tutorial]$ paste <(ls $d) <(perl -le 'sub p{my
$l=pop @_;unless(@_){return map [$_],@$l;}return map { my $ll=$_; map
[@$ll,$_],@$l} p(@_);} @a=[A,C,G,T]; print join("", @$_) for
p(@a,@a,@a,@a,@a,@a,@a,@a);' | awk -v k=$t 'NR<=k{print}') | awk
'BEGIN{print "sample-id\tbarcode-sequence\n#q2:types\tcategorical"}1'
> sample_metadata.tsv
[User@HPC ~/user/qiime2_tutorial]$ cat sample_metadata.tsv
sample-id    barcode-sequence
#q2:types    categorical
109-2    AAAAAAAA
1-1    AAAAAAAC
110-2    AAAAAAAG
113-2    AAAAAAAT
114-2    AAAAAACA
115-2    AAAAAACC
117-2    AAAAAACG
119-2    AAAAAACT
120-2    AAAAAAGA
126-2    AAAAAAGC
128-2    AAAAAAGG
130-2    AAAAAAGT
13-1    AAAAAATA
132-2    AAAAAATC
20-1    AAAAAATG
27-1    AAAAAATT
32-1    AAAAACAA
38-1    AAAAACAC
45-1    AAAAACAG
51-1    AAAAACAT
```

```
56-1    AAAAACCA
62-1    AAAAACCC
68-1    AAAAACCG
7-1    AAAAACCT
```

**Step 2**: Generate barcodes for each read using the file as above

```
[User@HPC ~/user/qiime2_tutorial]$ (for i in $(ls $d); do bc=$(awk -v
k=$i '$1==k{print $2}' sample_metadata.tsv); bioawk -cfastx -v k=$bc
'{print "@"$1" "$4"\n"k"\n+";for(i=0;i< length(k);i++){printf
"#"};printf "\n"}' $d/$i/Raw/*_R1_*.fastq ; done) > barcodes.fastq
```

Essentially, we are extracting the read headers from all the forward FASTQ files, and we assign the barcodes generated from sample_metadata.tsv file to those headers

```
[User@HPC ~/user/qiime2_tutorial]$ head barcodes.fastq
@M01359:18:000000000-A5HVT:1:1101:15648:3435 1:N:0:109
AAAAAAAA
+
########
@M01359:18:000000000-A5HVT:1:1101:15642:3453 1:N:0:109
AAAAAAAA
+
########
@M01359:18:000000000-A5HVT:1:1101:22693:3963 1:N:0:109
AAAAAAAA
[User@HPC ~/user/qiime2_tutorial]$ tail barcodes.fastq
+
########
@M01359:18:000000000-A5HVT:1:2114:15029:28668 1:N:0:68
AAAAACCG
+
########
@M01359:18:000000000-A5HVT:1:1106:13378:23600 1:N:0:7
AAAAACCT
+
########
```

**Step 3**: Collate all the forward reads from all the folders together in a single forward.fastq file

```
[User@HPC ~/user/qiime2_tutorial]$ (for i in $(ls $d); do cat
$d/$i/Raw/*_R1_*.fastq ; done) > forward.fastq
```

**Step 4**: Assemble all the reverse reads from all the folders together in a single reverse.fastq file

```
[User@HPC ~/user/qiime2_tutorial]$ (for i in $(ls $d); do cat
$d/$i/Raw/*_R2_*.fastq ; done) > reverse.fastq
```

See if the numbers match

```
[User@HPC ~/user/qiime2_tutorial]$ ls
barcodes.fastq  forward.fastq  reverse.fastq  sample_metadata.tsv
[User@HPC ~/user/qiime2_tutorial]$ bioawk -cfastx 'END{print NR}'
forward.fastq
554815
[User@HPC ~/user/qiime2_tutorial]$ bioawk -cfastx 'END{print NR}'
reverse.fastq
554815
[User@HPC ~/user/qiime2_tutorial]$ bioawk -cfastx 'END{print NR}'
barcodes.fastq
554815
[User@HPC ~/user/qiime2_tutorial]$
```

**Step 5**: Zip all the FASTQ files and move them to emp-paired-end-sequences folder

```
[User@HPC ~/user/qiime2_tutorial]$ gzip *.fastq
[User@HPC ~/user/qiime2_tutorial]$ mkdir emp-paired-end-sequences; mv
*.gz emp-paired-end-sequences/.
[User@HPC ~/user/qiime2_tutorial]$ ls
emp-paired-end-sequences  sample_metadata.tsv
[User@HPC ~/user/qiime2_tutorial]$
```

Next, we enable Qiime2 on the Orion cluster

```
[User@HPC ~/user/qiime2_tutorial]$ export
PATH=/home/opt/miniconda2/bin:$PATH
[User@HPC ~/user/qiime2_tutorial]$ source activate qiime2-2019.7
```

**Step 6**: Import the sequences to Qiime2

```
[User@HPC ~/user/qiime2_tutorial]$ qiime tools import --type
EMPPairedEndSequences --input-path emp-paired-end-sequences --output-
path emp-paired-end-sequences.qza
```

**Step 7**: Demultiplex the sequences in Qiime2

(qiime2-2019.7) [User@HPC ~/user/qiime2_tutorial]$ **qiime demux emp-paired --p-no-golay-error-correction --i-seqs emp-paired-end-sequences.qza --m-barcodes-file sample_metadata.tsv --m-barcodes-column barcode-sequence --o-per-sample-sequences demux.qza --o-error-correction-details demux-details.qza**

**Step 8**: Depends on the quality of your run, we want to fine tune Dada2 algorithm by specifying the thresholds

(qiime2-2019.7) [User@HPC ~/user/qiime2_tutorial]$ **qiime demux summarize --i-data ./demux.qza --o-visualization ./demux.qzv**

(qiime2-2019.7) [User@HPC ~/user/qiime2_tutorial]$ **qiime tools export --input-path demux.qzv --output-path output**

Next download the file to your local computer

**scp:~/user/qiime2_tutorial/demux.qzv .**

Next drag and drop the file on the Qiime2 viewer https://view.qiime2.org and manually figure out the thresholds, i.e., where the quality drops down significantly

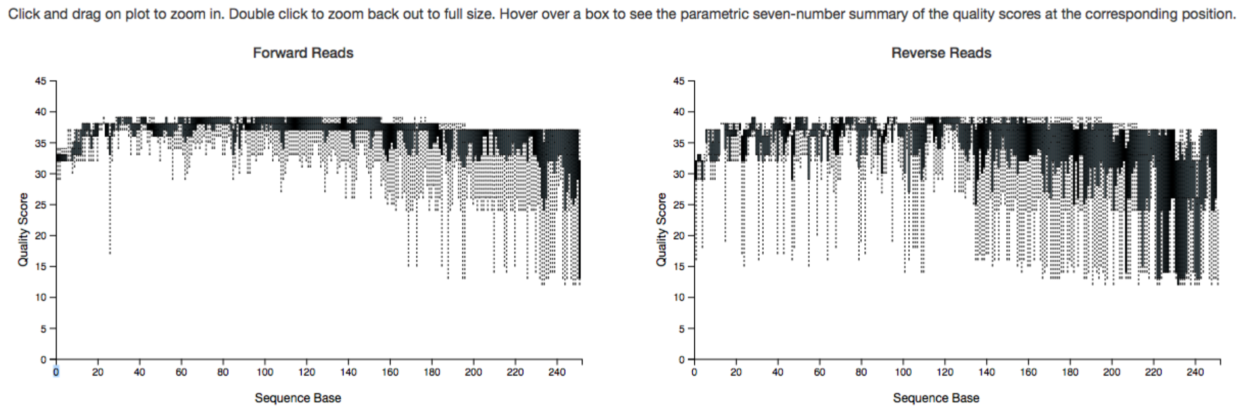


**Step 9**: Run Dada2 algorithm

(qiime2-2019.7) [User@HPC ~/user/qiime2_tutorial]$ **qiime dada2 denoise-paired --i-demultiplexed-seqs demux.qza --p-trim-left-f 0 --p-trim-left-r 0 --p-trunc-len-f 240 --p-trunc-len-r 200 --p-n-threads 0 --o-table table.qza --o-representative-sequences rep-seqs.qza --o-denoising-stats denoising-stats.qza --verbose**

**Step 10**: Generate the phylogenetic tree for the ASVs

```
(qiime2-2019.7) [User@HPC ~/user/qiime2_tutorial]$ unset
MAFFT_BINARIES
(qiime2-2019.7) [User@HPC ~/user/qiime2_tutorial]$ qiime phylogeny
align-to-tree-mafft-fasttree --i-sequences rep-seqs.qza --o-alignment
aligned-rep-seqs.qza --o-masked-alignment masked-aligned-rep-seqs.qza
--p-n-threads 0 --o-tree unrooted-tree.qza --o-rooted-tree rooted-
tree.qza
```

**Step 11**: Generate taxonomy for these ASVs

Using either SILVA 138

```
(qiime2-2019.7) [User@HPC ~/user/qiime2_tutorial]$ qiime feature-
classifier classify-sklearn --i-classifier ~/SILVA138_Database/silva-
138-99-nb-classifier.qza --i-reads rep-seqs.qza --o-classification
taxonomy.qza
```

Or MiDAS 3.7

```
(qiime2-2019.7) [User@HPC ~/user/qiime2_tutorial]$ qiime feature-
classifier classify-sklearn --i-classifier
/shared5/AD_Fanatics/MIDAS_TOUCH/Midas-classifier.qza --i-reads rep-
seqs.qza --o-classification taxonomy_MIDAS.qza
```

**Step 12**: Export all the files that Qiime2 generated

```
(qiime2-2019.7) [User@HPC ~/user/qiime2_tutorial]$ qiime tools export
--input-path table.qza --output-path output
Exported table.qza as BIOMV210DirFmt to directory output
(qiime2-2019.7) [User@HPC ~/user/qiime2_tutorial]$ qiime tools export
--input-path rep-seqs.qza --output-path output
Exported rep-seqs.qza as DNASequencesDirectoryFormat to directory
output
(qiime2-2019.7) [User@HPC ~/user/qiime2_tutorial]$ qiime tools export
--input-path rooted-tree.qza --output-path output

Exported rooted-tree.qza as NewickDirectoryFormat to directory output
(qiime2-2019.7) [User@HPC ~/user/qiime2_tutorial]$ qiime tools export
--input-path taxonomy.qza --output-path output

Exported taxonomy.qza as TSVTaxonomyDirectoryFormat to directory
output
```

```
(qiime2-2019.7) [User@HPC ~/user/qiime2_tutorial/output]$ ls
data.jsonp                          overview.html
demultiplex-summary.pdf             per-sample-fastq-counts.csv
demultiplex-summary.png             q2templateassets
dist                                quality-plot.html
dna-sequences.fasta                 reverse-seven-number-summaries.csv
feature-table.biom                  taxonomy.tsv
forward-seven-number-summaries.csv  tree.nwk
index.html
(qiime2-2019.7) [User@HPC ~/user/qiime2_tutorial/output]$
```

**Step 13**: Attach the abundance table of ASVs with their corresponding taxonomy to generate the biom file that you will use in the downstream statistical analysis. **For making Biome file compatible with R and phyloseq**, please check https://github.com/joey711/phyloseq/issues/821. Go inside the "output" folder generated in the previous step and write these commands

```
(qiime2-2019.7) [User@HPC ~/user/qiime2_tutorial/output]$ biom convert
-i feature-table.biom -o feature-table.tsv --to-tsv
(qiime2-2019.7) [User@HPC ~/user/qiime2_tutorial/output]$ head
feature-table.tsv
# Constructed from biom file
#OTU ID    1-1    109-2    110-2    113-2    114-2    115-2    117-2
119-2    120-2    126-2    128-2    13-1    130-2    132-2    20-1
27-1    32-1    38-1    45-1    51-1    56-1    62-1    68-1
5a6c87d6a4eb5e114959f6192f29b641    56.0    0.0    118.0    0.0
169.0    82.0    54509.0    452.0    0.0    0.0    1990.0    639.0
110.0    0.0    41.0    0.0    271.0    0.0    0.0    0.0    0.0
0.0    0.0
c6c3ab4e828fb40d6e05967b7aac9338    4090.0    227.0    67.0    0.0
587.0    29.0    2706.0    563.0    185.0    107.0    479.0    1015.0
191.0    819.0    62.0    0.0    0.0    0.0    684.0    3684.0
7426.0    247.0    6.0
35cbb315e019e0a8768d4b73862df415    579.0    89.0    352.0    0.0
939.0    459.0    10921.0    907.0    538.0    201.0    1386.0
3024.0    184.0    185.0    82.0    0.0    466.0    0.0    0.0
631.0    0.0    0.0    0.0
35ffcc3b809d667286737d79670b8de5    886.0    423.0    234.0    105.0
183.0    175.0    4576.0    211.0    604.0    170.0    654.0    1446.0
309.0    98.0    0.0    86.0    77.0    0.0    0.0    907.0    128.0
29.0    543.0
c24e0e391aa836b5eae25567c7eb89ee    80.0    74.0    15.0    0.0
14.0    0.0    90.0    0.0    17.0    0.0    0.0    182.0    27.0
```

```
0.0	20.0	4190.0	0.0	1181.0	130.0	19.0	4257.0
69.0	127.0
fd496fd32dc8c08ade2e8b6c9d8ee13d	119.0	59.0	10.0	11.0
0.0	0.0	3122.0	96.0	178.0	8.0	0.0	150.0	36.0
43.0	6185.0	0.0	0.0	0.0	0.0	119.0	0.0	10.0
172.0
2c5e87b291147c04e1b8d1c808b1aee0	165.0	0.0	67.0	0.0
117.0	0.0	0.0	0.0	0.0	0.0	987.0	3141.0	0.0
0.0	114.0	0.0	169.0	0.0	0.0	0.0	5030.0	0.0
0.0
9639a3291729a3758207b47715d9205f	0.0	43.0	149.0	0.0
178.0	0.0	6321.0	321.0	94.0	59.0	343.0	0.0
23.0	1336.0	0.0	0.0	52.0	0.0	12.0	201.0	0.0
0.0	0.0
(qiime2-2019.7) [User@HPC ~/user/qiime2_tutorial/output]$
```

```
(qiime2-2019.7) [User@HPC ~/user/qiime2_tutorial/output]$ sed -i s/Taxon/taxonomy/ taxonomy.tsv | sed -i s/Feature\ ID/FeatureID/ taxonomy.tsv
```

```
(qiime2-2019.7) [User@HPC ~/user/qiime2_tutorial/output]$ biom add-metadata  -i feature-table.tsv  -o feature_w_tax.biom  --observation-metadata-fp taxonomy.tsv  --observation-header FeatureID,taxonomy,Confidence  --sc-separated taxonomy --float-fields Confidence
```

### Bayesian Lowest Common Ancestor Algorithm (as an alternative to Naive-Bayes classifier)

Bayesian LCA-based Taxonomic Classification Method (BLCA) is a Bayesian-based method that provides a solid probabilistic basis for evaluating the taxonomic assignments for the query sequences with bootstrap confidence scores, which is based on Bayesian posterior probability that quantitatively weigh each database hit sequence according to its similarity to the query sequence - the more similar database hit sequence to the query, the more its contribution to the taxonomic assignment of the query. The relevant software to do this is available at https://github.com/qunfengdong/BLCA

### BLCA approach for SILVA138/MIDAS

We index the database to be used with BLCA

```
(qiime2-2019.7) [uzi@HPC ~/SILVA138_Database]$ makeblastdb -in dna-sequences.fasta -dbtype nucl -parse_seqids -out dna-sequences.fasta


Building a new DB, current time: 07/08/2020 10:23:39
New DB name:   ~/SILVA138_Database/dna-sequences.fasta
New DB title:  dna-sequences.fasta
Sequence type: Nucleotide
Keep MBits: T
Maximum file size: 1000000000B
Adding sequences from FASTA; added 436680 sequences in 19.3583 seconds.
```

The taxonomy that was exported using `qiime tools export` command is saved as taxonomy.tsv file, we remove the first line and save it. Afterwards, we format it

```
(qiime2-2019.7) [uzi@HPC ~/SILVA138_Database]$ tr "; " ";\t" < taxonomy.tsv > taxonomy.txt
```

```
(qiime2-2019.7) [uzi@HPC ~/SILVA138_Database]$ awk -F"\t" '{a=$1"\tspecies:"$8";genus:"$7";family:"$6";order:"$5";class:"$4";phylum:"$3";superkingdom:"$2; gsub(";;",";",a);gsub(";$","",a); gsub(":s__|:g__|:f__|:o__|:c__|:p__|:d__",":",a);print a}' taxonomy.txt > taxonomy_BLCA.txt
```

**Step 1**: Run BLCA

Option 1 (SILVA138):

```
(python3) [User@HPC ~/BLCA]$  2.blca_main.py -i dna-sequences.fasta -q ~/SILVA138_Database/dna-sequences.fasta -r ~SILVA138_Database/taxonomy_BLCA.txt
```

Option 2 (MIDAS 3.7):

```
(python3) [User@HPC ~/BLCA]$  2.blca_main.py -i dna-sequences.fasta -q ~/MIDAS3.7_Database/Midas.fasta -r ~/MIDAS3.7_Database /Midas_taxonomy_BLCA.txt
```

clustalo is located in your PATH!

> > Fasta file read in!

> > Reading in taxonomy information! ....

blastn is located in your PATH!

> > Running blast!!

> > Blastn Finished!!

> > read in blast file...

> > blastn file opened

> > blast output read in

> > Start aligning reads...

> > Taxonomy file generated!!

Time elapsed: 22 minutes

**Step 2**: Convert the BLCA format to regular qiime2 taxonomy format

```
(python3) [User@HPC ~/BLCA]$ awk -F"\t" 'BEGIN{print
"FeatureID\tTaxon\tConfidence"}{gsub("superkingdom:","D_0__",$2);gsub("phy
lum:","D_1__",$2);gsub("class:","D_2__",$2);gsub("order:","D_3__",$2);gsub
("family:","D_4__",$2);gsub("genus:","D_5__",$2);gsub("species:","D_6__",$
2);gsub(";$|\t$","",$2);gsub(";"," ;",$2);gsub(" ;[0-9]+(\.[0-9]+)?
;",";",$2);gsub("Unclassified","Unassigned\t100.0",$2);gsub("
+;","\t",$2);print $1"\t"$2}' dna-sequences.fasta.blca.out >
taxonomy_midas_BLCA.tsv
```

**Step 3**: Create the biom file

```
 (qiime2-2019.7) [User@HPC ~/BLCA]$ biom add-metadata  -i feature-
table.tsv  -o feature_w_tax_midas_BLCA.biom  --observation-metadata-fp
taxonomy_midas_BLCA.tsv  --observation-header
FeatureID,taxonomy,Confidence  --sc-separated taxonomy --float-fields
Confidence
```

### Collating multiple V regions and generating a collated phylogenetic tree

**Step 1**: We first need to put all the biom files in a “data” subdirectory and run the following two scripts in R:

collated_biom_part1.R script:

```
#From the RStudio menu, click on "Session" and then "Set Working
Directory" to "To Source File Location"
#Collation script to combine biom tables from multiple regions (Part 1
to assemble OTUs/ASVs at species level)
```

```
library(phyloseq)
library(vegan)
library(ecodist)

#PARAMETERS ###########################
files <- list.files(path="data/",pattern = "biom") #Get the list of
biom files in subfolder data
#PARAMETERS ###########################


#Iterate through all the biom files
for(i in files){
  tmp<-import_biom(paste("data/",i,sep=""))
  tmp_Rank7<-tax_glom(tmp, "Rank7")
  ft<-as(t(otu_table(tmp)),"matrix")
  st<-as(t(otu_table(tmp_Rank7)),"matrix")
  #There was a bug where if you do not get species level assignment
then some of the samples are completely zero,
  #and downstream Bray-Curtis distance gives NaN value, resolution is
to get rid of those samples from both
  #st and ft
  st<-st[rowSums(st)>0,]
  ft<-ft[rownames(st),]
  full.dist<-vegdist(ft,method="bray")
  Rank7.dist<-vegdist(st,method="bray")

capture.output(mantel(full.dist~Rank7.dist,nperm=10000)[c("mantelr","p
val1")],file=paste("data/",gsub("\\.biom","",i),"_mantel.txt",sep=""))

capture.output(protest(monoMDS(full.dist),monoMDS(Rank7.dist),symmetri
c=T),file=paste("data/",gsub("\\.biom","",i),"_procrustes.txt",sep="")
)
  #Next see how much reads we have lost by getting rid of anything
that doesn't have species resolution
  stats<-
data.frame(all=rowSums(t(otu_table(tmp))),species=rowSums(t(otu_table(
tmp_Rank7))),archael_species=tryCatch(rowSums(t(otu_table(tmp_Rank7)[g
repl("Archaea",tax_table(tmp_Rank7)[,1]),])),error=function(e)
data.frame(row.names=rownames(t(otu_table(tmp_Rank7))),archael_species
=rep(0,nrow(t(otu_table(tmp_Rank7)))))),bacterial_species=rowSums(t(ot
u_table(tmp_Rank7)[grepl("Bacteria",tax_table(tmp_Rank7)[,1]),])))

write.csv(otu_table(tmp_Rank7),file=paste("data/",gsub("\\.biom","",i)
,"_abund_table_Rank7.csv",sep=""))
```

```
write.csv(tax_table(tmp_Rank7),file=paste("data/",gsub("\\.biom","",i)
,"_tax_table_Rank7.csv",sep=""))

write.csv(stats,paste("data/",gsub("\\.biom","",i),"_stats_Rank7.csv",
sep=""))
  cat(paste(i,"processed!\n"))
}
```

collated_biom_part2.R script:

```
#From the RStudio menu, click on "Session" and then "Set Working
Directory" to "To Source File Location"
#Collation script to combine biom tables from multiple regions (Part 2
to generate the biom file as well as taxonomy)

library(phyloseq)
library(biomformat)

#PARAMETERS ###########################
abund_files <- list.files(path="data/",pattern =
"abund_table_Rank7.csv") #Get the list of biom files in subfolder data
tax_files<-list.files(path="data/",pattern = "tax_table_Rank7.csv")
#Get the list of biom files in subfolder data
#PARAMETERS ###########################


#Get the file_prefixes
file_prefixes<-gsub("_abund_table_Rank7.csv","",abund_files)

#PASS 1 where we just extract the list of samples and taxa

#global list of all the taxa
taxa_global<-NULL
#global list of all the samples
samples_global<-NULL
for(i in file_prefixes){
  #extract the abundance table
  at<-
read.csv(paste("data/",i,"_abund_table_Rank7.csv",sep=""),header=T,row
.names=1,check.names=FALSE)
  #extract the taxonomy table
  tt<-
read.csv(paste("data/",i,"_tax_table_Rank7.csv",sep=""),header=T,row.n
ames=1,check.names=FALSE)
```

```
  #extract the list of unique taxa in the taxonomy table
  tmp<-as.character(unique(tt[,"Rank7"]))
  tmp2<-as.character(unique(colnames(at)))
  if(is.null(taxa_global)){taxa_global<-tmp} else {taxa_global<-
unique(c(taxa_global,tmp))}
  if(is.null(samples_global)){samples_global<-tmp2} else
{samples_global<-unique(c(samples_global,tmp2))}
}

#now that we have the complete list of samples and taxa, we need to
make a collated abundance table, as well as taxonomy table
collated_table<-
matrix(ncol=length(samples_global),nrow=length(taxa_global),data=0)
colnames(collated_table)<-samples_global
rownames(collated_table)<-taxa_global
collated_table<-as.data.frame(collated_table)
collated_taxonomy<-
matrix(ncol=7,nrow=length(taxa_global),data="empty")
colnames(collated_taxonomy)<-
c("Rank1","Rank2","Rank3","Rank4","Rank5","Rank6","Rank7")
rownames(collated_taxonomy)<-taxa_global
collated_taxonomy<-as.data.frame(collated_taxonomy)
collated_taxonomy[] <- lapply(collated_taxonomy, function(x)
as.character(x))

#PASS 2 where we populate collated_table and collated_taxonomy

for(i in file_prefixes){
  #extract the abundance table
  at<-
read.csv(paste("data/",i,"_abund_table_Rank7.csv",sep=""),header=T,row
.names=1,check.names=FALSE)
  #extract the taxonomy table
  tt<-
read.csv(paste("data/",i,"_tax_table_Rank7.csv",sep=""),header=T,row.n
ames=1,check.names=FALSE)

  #we extract at2 and give it taxonomy names
  at2<-at
  rownames(at2)<-paste("Taxa_",seq(1,nrow(at)),sep="")
  tt2<-tt
  rownames(tt2)<-paste("Taxa_",seq(1,nrow(at)),sep="")

  #Explicitly convert tt2 to character
  tt2[] <- lapply(tt2, function(x) as.character(x))
```

```
    for(j in rownames(at2)){
    for(k in colnames(at2) ){
      collated_table[tt2[j,"Rank7"],k]<-
collated_table[tt2[j,"Rank7"],k]+at2[j,k]
    }
  }
  for(j in rownames(tt2)){
    for(k in colnames(tt2)){
      collated_taxonomy[tt2[j,"Rank7"],k]<-tt2[j,k]
    }
  }
}

#Now we assign Taxa_1, Taxa_2 to both collated_table, and
collated_taxonomy
rownames(collated_table)<-
paste("Taxa_",seq(1,nrow(collated_table)),sep="")
rownames(collated_taxonomy)<-
paste("Taxa_",seq(1,nrow(collated_taxonomy)),sep="")

#We construct a biom object
suppressWarnings(ps.b <- biomformat::make_biom(
  data=collated_table,
  id="No Table ID",
  sample_metadata=NULL,
  observation_metadata=collated_taxonomy,
  matrix_element_type="int")
)

#export the biom file
biomformat::write_biom(x = ps.b, biom_file =
"collated_feature_w_tax.biom")
write.csv(collated_taxonomy,file="collated_taxonomy.csv")

# In the scripts where you use phyloseq, you have two options to
# load the biom file (both will generate the physeq object). The
# reason why we have these two options is mainly because write_biom()
# from biomformat package saves biom file in a format that is not
readable
# using import_biom() function. So we are introducing an extra step:
#
# physeq<-import_biom("collated_feature_w_tax.biom")
# b_<-read_biom("collated_feature_w_tax.biom");physeq<-
merge_phyloseq(otu_table(as(biom_data(b_),"matrix"),taxa_are_rows=TRUE
),tax_table(as(observation_metadata(b_),"matrix")))
```

After running `collated_biom_part1.R` and `collated_biom_part2.R` to generate `collated_taxonomy.csv`, we will use the following commands to generate a collated phylogenetic tree.

**Step 2**: Upload collated_taxonomy.csv to the server from your local folder

```
scp collated_taxonomy.csv:
~/collation/collated_taxonomy_midas_BLCA.csv
```

It has the following format:

```
[User@HPC ~/collation]$ head collated_taxonomy_midas_BLCA.csv
"","Rank1","Rank2","Rank3","Rank4","Rank5","Rank6","Rank7"
"Taxa_1","D_0__Bacteria","D_1__Bacteroidetes","D_2__Bacteroidia","D_3_
_Sphingobacteriales","D_4__Lentimicrobiaceae","D_5__midas_g_362","D_6_
_midas_s_5396"
"Taxa_2","D_0__Bacteria","D_1__Firmicutes","D_2__Clostridia","D_3__Clo
stridiales","D_4__Eubacteriaceae","D_5__midas_g_2229","D_6__midas_s_22
29"
"Taxa_3","D_0__Bacteria","D_1__Firmicutes","D_2__Bacilli","D_3__Lactob
acillales","D_4__Streptococcaceae","D_5__Lactococcus","D_6__Lactococcu
s_chungangensis"
"Taxa_4","D_0__Bacteria","D_1__Firmicutes","D_2__Clostridia","D_3__Clo
stridiales","D_4__Eubacteriaceae","D_5__midas_g_2229","D_6__midas_s_33
25"
"Taxa_5","D_0__Bacteria","D_1__Firmicutes","D_2__Clostridia","D_3__Clo
stridiales","D_4__Ruminococcaceae","D_5__Ruminococcaceae_UCG-
014","D_6__midas_s_4482"
"Taxa_6","D_0__Bacteria","D_1__Bacteroidetes","D_2__Bacteroidia","D_3_
_Bacteroidales","D_4__Paludibacteraceae","D_5__Paludibacter","D_6__mid
as_s_2705"
"Taxa_7","D_0__Bacteria","D_1__Firmicutes","D_2__Bacilli","D_3__Lactob
acillales","D_4__midas_f_4","D_5__midas_g_4853","D_6__midas_s_4853"
"Taxa_8","D_0__Bacteria","D_1__Proteobacteria","D_2__Gammaproteobacter
ia","D_3__Pseudomonadales","D_4__Pseudomonadaceae","D_5__Pseudomonas",
"D_6__Pseudomonas_caeni"
"Taxa_9","D_0__Bacteria","D_1__Proteobacteria","D_2__Gammaproteobacter
ia","D_3__Enterobacteriales","D_4__Enterobacteriaceae","D_5__Enterobac
ter","D_6__midas_s_9574"
```

**Step 3**: Remove inverted commas, get rid of the first line from the taxonomy file, and convert comma-delimited to tab-delimited, and fix "species:" bug that arises in MIDAS, and convert to original MIDAS taxonomy format and also remove tabs from after kingdom assignment

```
[User@HPC ~/collation]$ sed 's/"//g' collated_taxonomy_midas_BLCA.csv
| tr "," "\t" | sed 1d | sed 's/species:/D_6__/g' | awk
'{gsub("D_0__","k__",$0);gsub("D_1__","p__",$0);gsub("D_2__","c__",$0)
;gsub("D_3__","o__",$0);gsub("D_4__","f__",$0);gsub("D_5__","g__",$0);
gsub("D_6__","s__",$0);gsub("\tp__","; p__",$0);gsub("\tc__",";
c__",$0);gsub("\tp__","; p__",$0);gsub("\to__",";
o__",$0);gsub("\tp__","; p__",$0);gsub("\tf__",";
f__",$0);gsub("\tg__","; g__",$0);gsub("\ts__","; s__",$0)}1' >
collated_taxonomy_midas_BLCA.tsv
```

Sanity check: see if the format is exactly the same as the MIDAS database

```
[User@HPC ~/collation]$ head collated_taxonomy_midas_BLCA.tsv
Taxa_1    k__Bacteria; p__Bacteroidetes; c__Bacteroidia;
o__Sphingobacteriales; f__Lentimicrobiaceae; g__midas_g_362;
s__midas_s_5396
Taxa_2    k__Bacteria; p__Firmicutes; c__Clostridia; o__Clostridiales;
f__Eubacteriaceae; g__midas_g_2229; s__midas_s_2229
Taxa_3    k__Bacteria; p__Firmicutes; c__Bacilli; o__Lactobacillales;
f__Streptococcaceae; g__Lactococcus; s__Lactococcus_chungangensis
Taxa_4    k__Bacteria; p__Firmicutes; c__Clostridia; o__Clostridiales;
f__Eubacteriaceae; g__midas_g_2229; s__midas_s_3325
Taxa_5    k__Bacteria; p__Firmicutes; c__Clostridia; o__Clostridiales;
f__Ruminococcaceae; g__Ruminococcaceae_UCG-014; s__midas_s_4482
Taxa_6    k__Bacteria; p__Bacteroidetes; c__Bacteroidia;
o__Bacteroidales; f__Paludibacteraceae; g__Paludibacter;
s__midas_s_2705
Taxa_7    k__Bacteria; p__Firmicutes; c__Bacilli; o__Lactobacillales;
f__midas_f_4; g__midas_g_4853; s__midas_s_4853
Taxa_8    k__Bacteria; p__Proteobacteria; c__Gammaproteobacteria;
o__Pseudomonadales; f__Pseudomonadaceae; g__Pseudomonas;
s__Pseudomonas_caeni
Taxa_9    k__Bacteria; p__Proteobacteria; c__Gammaproteobacteria;
o__Enterobacteriales; f__Enterobacteriaceae; g__Enterobacter;
s__midas_s_9574
Taxa_10    k__Bacteria; p__Proteobacteria; c__Gammaproteobacteria;
o__Enterobacteriales; f__Enterobacteriaceae; g__Escherichia-Shigella;
s__midas_s_9539
```

```
[User@HPC ~/collation]$ head ~/MIDAS3.7_Database /Midas_taxonomy.txt
```

```
FLASV1.1417    k__Bacteria; p__Chloroflexi; c__Anaerolineae;
o__Caldilineales; f__Amarolineaceae; g__Ca_Amarolinea; s__midas_s_1
FLASV2.1445    k__Bacteria; p__Actinobacteria; c__Acidimicrobiia;
o__Microtrichales; f__Microtrichaceae; g__Ca_Microthrix; s__midas_s_2
FLASV3.1527    k__Bacteria; p__Actinobacteria; c__Acidimicrobiia;
o__Microtrichales; f__Microtrichaceae; g__Ca_Microthrix;
s__Ca_Microthrix_parvicella
FLASV4.1481    k__Bacteria; p__Firmicutes; c__Bacilli;
o__Lactobacillales; f__Carnobacteriaceae; g__Trichococcus;
s__midas_s_4
FLASV5.1442    k__Bacteria; p__Actinobacteria; c__Actinobacteria;
o__Micrococcales; f__Intrasporangiaceae; g__Tetrasphaera; s__midas_s_5
FLASV6.1456    k__Bacteria; p__Bacteroidetes; c__Bacteroidia;
o__Chitinophagales; f__Saprospiraceae; g__midas_g_6; s__midas_s_6
FLASV7.1445    k__Bacteria; p__Firmicutes; c__Clostridia;
o__Clostridiales; f__Syntrophomonadaceae; g__midas_g_7; s__midas_s_7
FLASV8.1456    k__Bacteria; p__Firmicutes; c__Clostridia;
o__Clostridiales; f__Ruminococcaceae; g__Fastidiosipila; s__midas_s_8
FLASV9.1457    k__Bacteria; p__Firmicutes; c__Clostridia;
o__Clostridiales; f__Ruminococcaceae; g__Fastidiosipila; s__midas_s_8
FLASV10.1442    k__Bacteria; p__Actinobacteria; c__Actinobacteria;
o__Micrococcales; f__Intrasporangiaceae; g__Tetrasphaera; s__midas_s_5
```

**PASS 1**: Those that are returned with same taxonomy as MIDAS database, we will use this to replace first column with well resolved IDs

**MIDAS:**

```
[User@HPC ~/collation]$ awk -F"\t" 'BEGIN{while((getline k
<"~/MIDAS3.7_Database/Midas_taxonomy.txt")>0){split(k,a,"\t");i[a[2]]=
a[1]}} {if(i[$2]){ $1=i[$2]}}1' collated_taxonomy_midas_BLCA.tsv >
collated_taxonomy_midas_BLCA_level1.tsv
```

**SILVA138:**

```
[User@HPC ~/collation]$ awk -F"\t" 'BEGIN{while((getline k
<"~/SILVA138_Database/silva-138-99-
tax/taxonomy.tsv")>0){split(k,a,"\t");i[a[2]]=a[1]}} {if(i[$2]){
$1=i[$2]}}1' collated_taxonomy.tsv > collated_taxonomy_level1.tsv
```

Sanity Check: See how many taxa are not assigned

```
[User@HPC ~/collation]$ tail collated_taxonomy_midas_BLCA_level1.tsv
```

```
FLASV9428.1460 k__Bacteria; p__Proteobacteria; c__Gammaproteobacteria;
o__Betaproteobacteriales; f__Rhodocyclaceae; g__Thauera;
s__Thauera_terpenica
FLASV7402.1447 k__Bacteria; p__Bacteroidetes; c__Bacteroidia;
o__Bacteroidales; f__Bacteroidetes_vadinHA17; g__midas_g_19;
s__midas_s_2210
FLASV7071.1462 k__Bacteria; p__Proteobacteria; c__Gammaproteobacteria;
o__Betaproteobacteriales; f__Rhodocyclaceae; g__Ca_Accumulibacter;
s__midas_s_168
FLASV8700.1425 k__Bacteria; p__Chloroflexi; c__Anaerolineae;
o__Anaerolineales; f__Anaerolineaceae; g__midas_g_156; s__midas_s_876
FLASV6843.1458 k__Bacteria; p__Proteobacteria; c__Gammaproteobacteria;
o__Betaproteobacteriales; f__Rhodocyclaceae; g__Dechloromonas;
s__midas_s_96
FLASV1294.1462 k__Bacteria; p__Proteobacteria; c__Gammaproteobacteria;
o__Betaproteobacteriales; f__Rhodocyclaceae; g__Azospira;
s__midas_s_1294
FLASV6109.1507 k__Bacteria; p__Atribacteria; c__JS1; o__midas_o_230;
f__midas_f_230; g__midas_g_6109; s__midas_s_6109
FLASV4168.1451 k__Bacteria; p__Proteobacteria; c__Gammaproteobacteria;
o__Betaproteobacteriales; f__Burkholderiaceae; g__midas_g_81;
s__midas_s_4168
Taxa_2698    k__Bacteria; p__Proteobacteria; c__Gammaproteobacteria;
o__Betaproteobacteriales; f__Burkholderiaceae; g__Diaphorobacter;
s__midas_s_7296
FLASV9485.1454 k__Bacteria; p__Firmicutes; c__Clostridia;
o__Clostridiales; f__Lachnospiraceae; g__Fusicatenibacter;
s__Fusicatenibacter_saccharivorans
```

**PASS 2**: We rerun everything on level1 file, but instead of matching the whole taxonomy, we match the last taxonomic level

MIDAS 3.7:

```
[User@HPC ~/collation]$ awk -F"\t" 'BEGIN{while((getline k
<"~/MIDAS3.7_Database/Midas_taxonomy.txt")>0){split(k,a,"\t");split(a[
2],b,"; ");i[b[7]]=a[1]}} {if($1~/Taxa_/){ split($2,b,";
");$1=i[b[7]]}}1' collated_taxonomy_midas_BLCA_level1.tsv >
collated_taxonomy_midas_BLCA_level2.tsv
```

SILVA138:

```
[User@HPC ~/collation]$ awk -F"\t" 'BEGIN{while((getline k
<"~/SILVA138_Database/silva-138-99-
```

```
tax/taxonomy.tsv")>0){split(k,a,"\t");split(a[2],b,";
");i[b[7]]=a[1]}} {if($1~/Taxa_/){ split($2,b,"; ");$1=i[b[7]]}}1'
collated_taxonomy_level1.tsv > collated_taxonomy_level2.tsv
```

Sanity Check: See if it has worked

```
[User@HPC ~/collation]$ tail collated_taxonomy_midas_BLCA_level2.tsv
FLASV9428.1460 k__Bacteria; p__Proteobacteria; c__Gammaproteobacteria;
o__Betaproteobacteriales; f__Rhodocyclaceae; g__Thauera;
s__Thauera_terpenica
FLASV7402.1447 k__Bacteria; p__Bacteroidetes; c__Bacteroidia;
o__Bacteroidales; f__Bacteroidetes_vadinHA17; g__midas_g_19;
s__midas_s_2210
FLASV7071.1462 k__Bacteria; p__Proteobacteria; c__Gammaproteobacteria;
o__Betaproteobacteriales; f__Rhodocyclaceae; g__Ca_Accumulibacter;
s__midas_s_168
FLASV8700.1425 k__Bacteria; p__Chloroflexi; c__Anaerolineae;
o__Anaerolineales; f__Anaerolineaceae; g__midas_g_156; s__midas_s_876
FLASV6843.1458 k__Bacteria; p__Proteobacteria; c__Gammaproteobacteria;
o__Betaproteobacteriales; f__Rhodocyclaceae; g__Dechloromonas;
s__midas_s_96
FLASV1294.1462 k__Bacteria; p__Proteobacteria; c__Gammaproteobacteria;
o__Betaproteobacteriales; f__Rhodocyclaceae; g__Azospira;
s__midas_s_1294
FLASV6109.1507 k__Bacteria; p__Atribacteria; c__JS1; o__midas_o_230;
f__midas_f_230; g__midas_g_6109; s__midas_s_6109
FLASV4168.1451 k__Bacteria; p__Proteobacteria; c__Gammaproteobacteria;
o__Betaproteobacteriales; f__Burkholderiaceae; g__midas_g_81;
s__midas_s_4168
FLASV7296.1451 k__Bacteria; p__Proteobacteria; c__Gammaproteobacteria;
o__Betaproteobacteriales; f__Burkholderiaceae; g__Diaphorobacter;
s__midas_s_7296
FLASV9485.1454 k__Bacteria; p__Firmicutes; c__Clostridia;
o__Clostridiales; f__Lachnospiraceae; g__Fusicatenibacter;
s__Fusicatenibacter_saccharivorans
```

**Step 4**: Get all the reference IDs from the level 2 file just generated:

```
[User@HPC ~/collation]$ cut -d ' ' -f1
collated_taxonomy_midas_BLCA_level2.tsv > Reference_IDs.txt
```

Also, get the original Taxa numbers as well

```
[User@HPC ~/collation]$ awk -F"\t" '{print $1}'
collated_taxonomy_midas_BLCA.tsv > Taxa_IDs.txt
```

**Step 5**: Get all the sequences for these Reference_IDs.txt

MIDAS 3.7:

```
[User@HPC ~/collation]$ bioawk -cfastx 'BEGIN{while((getline k
<"Reference_IDs.txt")>0)i[k]=1}{if(i[$name])print ">"$name"\n"$seq}'
~/MIDAS3.7_Database/Midas.fasta > Taxa1.fa
```

SILVA138:

```
[User@HPC ~/collation]$ bioawk -cfastx 'BEGIN{while((getline k
<"Reference_IDs.txt")>0)i[k]=1}{if(i[$name])print ">"$name"\n"$seq}'
~/SILVA138_Database/silva-138-99-seqs/dna-sequences.fasta > Taxa1.fa
```

**Step 6**: Now generate the mapping file (MIDAS IDs to Taxa IDs in your sample)

```
[User@HPC ~/collation]$ paste Reference_IDs.txt Taxa_IDs.txt >
mapping.txt
```

**Step 7**: Replace the IDs with the actual Taxa numbers

```
[User@HPC ~/collation]$ bioawk -cfastx 'BEGIN{while((getline k
<"mapping.txt")>0){split(k,a,"\t");i[a[1]]=a[2]}}{print
">"i[$name]"\n"$seq}' Taxa1.fa > Taxa.fa
```

**Step 8:** Import the sequences to QIIME2

```
(qiime2-2019.7) [User@HPC ~/collation]$ qiime tools import --type
'FeatureData[Sequence]' --input-path Taxa.fa --output-path Taxa.qza
```

**Step 9**: Generate the phylogenetic tree

```
(qiime2-2019.7) [User@HPC ~/collation]$ unset MAFFT_BINARIES
(qiime2-2019.7) [User@HPC ~/collation]$ qiime phylogeny align-to-tree-
mafft-fasttree --i-sequences Taxa.qza --o-alignment aligned-rep-
seqs.qza --o-masked-alignment masked-aligned-rep-seqs.qza --p-n-
threads 0 --o-tree unrooted-tree.qza --o-rooted-tree rooted-tree.qza
```

**Step 10**: Export the phylogenetic tree to collated_taxonomy_midas_BLCA folder which contains tree.nwk file that you have to import back

```
qiime2-2019.7) [User@HPC ~/collation]$ qiime tools export --input-path
rooted-tree.qza --output-path collated_taxonomy_midas_BLCA
```
