## Supplementary Information File 3 for "Circular Economy of Anaerobic Biofilm Microbiomes: A Meta-Analysis Framework for Re-exploration of Amplicon Sequencing Data"

### Supplemental File 3 (Supplemental Figures)

**List of Figures**:

**Fig S1** – Bacterial and archaeal species-level resolution (i.e., all seven ranks were resolved) as a proportion of the total observed ASVs with Superkingdom tag as “Bacteria/Archaea” using both MiDAS 3.7 and SILVA138 databases where **(a)** and **(c)** use BLCA for bacteria and archaeal groups, respectively; **(b)** and **(d)** use NBC for bacterial and archaeal groups, respectively all comparing different v-regions.

**Fig S2** – Bacterial and archaeal species proportion with known nomenclature at “Species” rank i.e., doesn’t contain the pattern “Uncultured|uncultured|metagenome” for SILVA138, and “midas_s” for MIDAS 3.7 for a proportion of ASVs assigned to “Bacteria/Archaea” at superkingdom level where **(a)** and **(c)** use BLCA for bacteria and archaeal groups, respectively; **(b)** and **(d)** use NBC for bacterial and archaeal groups, respectively all comparing different v-regions.

**Fig S3** – Bacterial and archaeal species proportion with known nomenclature at “Species” rank i.e., doesn’t contain the pattern “Uncultured|uncultured|metagenome” for SILVA138, and “midas_s” for MIDAS 3.7 for a proportion of ASVs assigned to “Bacteria/Archaea” at superkingdom level where **(a)** and **(c)** use BLCA for bacteria and archaeal groups, respectively; **(b)** and **(d)** use NBC for bacterial and archaeal groups, respectively all comparing different extraction methods.

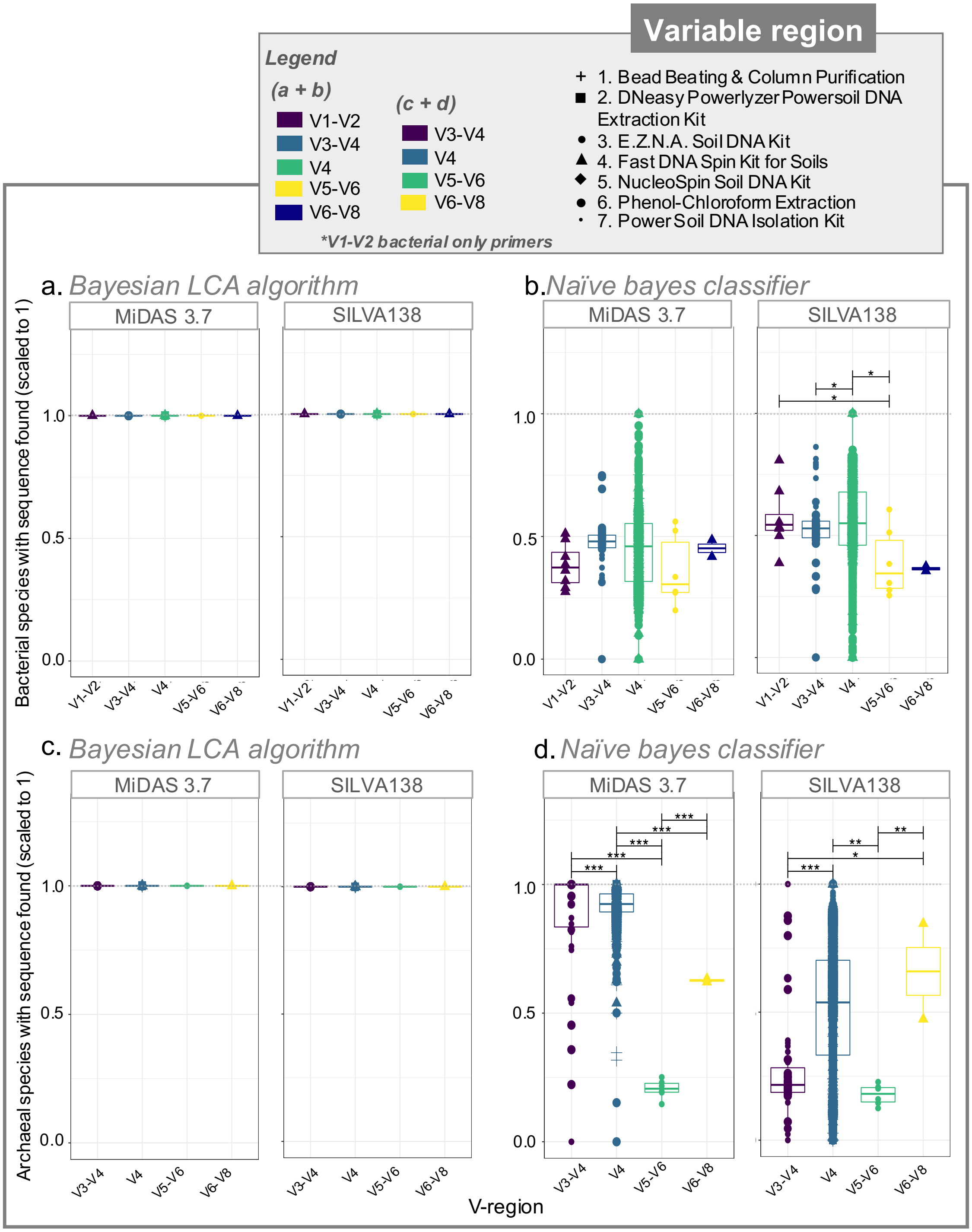


**Figure S1**. Bacterial and archaeal species-level resolution (i.e., all seven ranks were resolved) as a proportion of the total observed ASVs with Superkingdom tag as “Bacteria/Archaea” using both MiDAS 3.7 and SILVA138 databases where **(a)** and **(c)** use BLCA for bacteria and archaeal groups, respectively; **(b)** and **(d)** use NBC for bacterial and archaeal groups, respectively all comparing different v-regions.

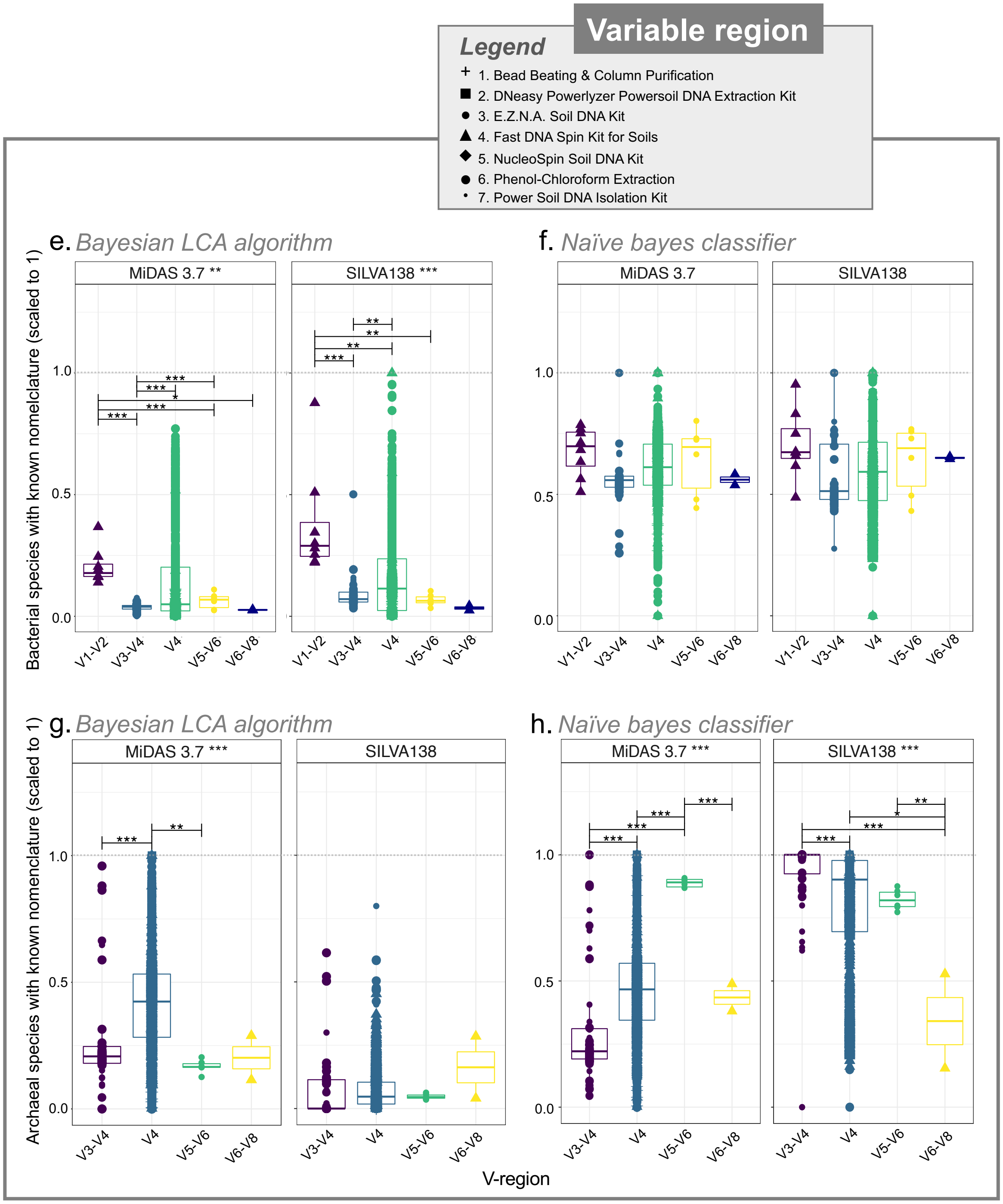


**Figure S2**. Bacterial and archaeal species proportion with known nomenclature at “Species” rank i.e., doesn’t contain the pattern “Uncultured|uncultured|metagenome” for SILVA138, and “midas_s” for MIDAS 3.7 for a proportion of ASVs assigned to “Bacteria/Archaea” at superkingdom level where **(a)** and **(c)** use BLCA for bacteria and archaeal groups, respectively; **(b)** and **(d)** use NBC for bacterial and archaeal groups, respectively all comparing different v-regions.

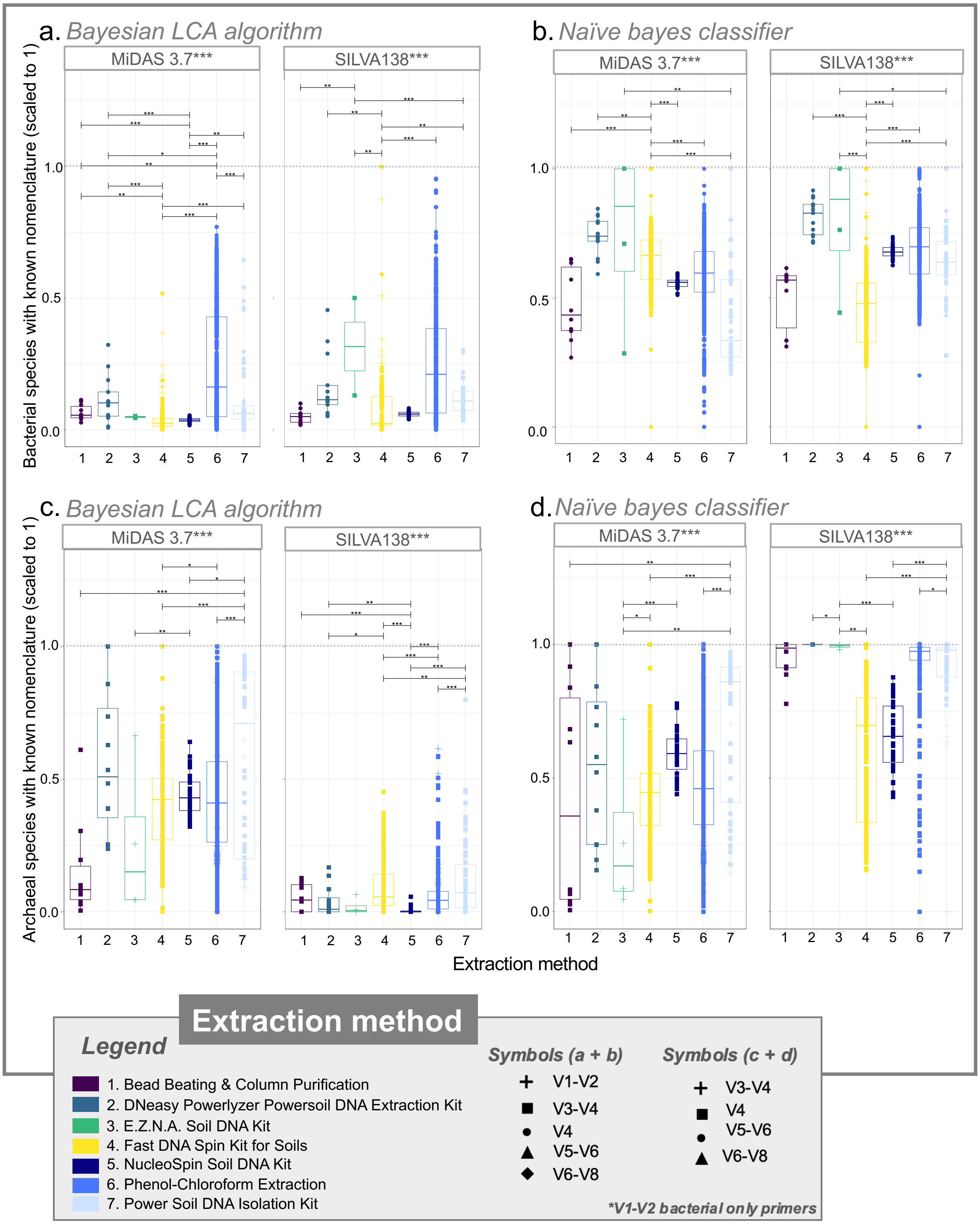


**Figure 4**. Bacterial and archaeal species proportion with known nomenclature at “Species” rank i.e., doesn’t contain the pattern “Uncultured|uncultured|metagenome” for SILVA138, and “midas_s” for MIDAS 3.7 for a proportion of ASVs assigned to “Bacteria/Archaea” at superkingdom level where **(a)** and **(c)** use BLCA for bacteria and archaeal groups, respectively;

**(b)** and **(d)** use NBC for bacterial and archaeal groups, respectively all comparing different extraction methods.
